# Di-codon organization links tRNA modifications to cancer cell proteome composition

**DOI:** 10.64898/2026.03.10.710911

**Authors:** Kiana H. Moghaddam, Clelia Timpone, Laasya Naga Gowda, Candani A. Tutuka, Gail P. Risbridger, Joel R. Steele, Ralf B. Schittenhelm, Eric P. Kusnadi, Luc Furic, Ola Larsson

## Abstract

Hormone-signalling modulates levels of tRNA-modifying enzymes, including ELP3, in cancer cells. Here we show that ELP3 is required to sustain proliferation in prostate cancer cells. Intriguingly, although ELP3 modifies tRNA at the U34-position, the ELP3-sensitive proteome was poorly explained by frequencies of codons requiring U34-modified tRNA for decoding. Instead, we identified six codon pairs (herein denoted ELP3 down di-codons “E3dDCs”) that, in concert with local sequence context and 5’UTR features affecting translation initiation, explain ELP3-sensitive protein expression. Moreover, despite activation of the canonical integrated stress response upon ELP3 suppression, the associated expression program was paradoxically suppressed in a fashion correlating with higher E3dDC frequencies. E3dDCs were also enriched in mitotic regulators, and ELP3-suppression caused mitotic defects that limited proliferation. Therefore, ELP3 regulates proteome composition via E3dDCs that, depending on their local sequence context and translation initiation activity, determine protein expression and downstream cellular phenotypes upon reduced levels of U34-modified tRNA.

## INTRODUCTION

tRNA modifications of the uridine at position 34 (U34) have well-established roles across the domains of life by ensuring optimal decoding fidelity during protein synthesis^1–6^. In eukaryotic cells, a cytosolic U34 modification pathway catalyzes the formation of 5-carboxymethyl (cm^5^)-based modifications on U34 via consecutive enzymatic activities of the Elongator (ELP) complex, the AlkB Homolog 8 tRNA Methyltransferase (ALKBH8) and cytosolic thiouridylases 1/2 (CTU1/2), giving rise to cm^5^, mcm^5^ and mcm^5^s^2^ modifications in a subset of tRNA^7,8^. Modifications at the U34 position play fundamental roles in maintaining cellular functions^8–12^. For example, defects in U34 tRNA modifications caused by mutations in Elongator subunits have been linked to neurodegenerative phenotypes^13,14^.

Intriguingly, in neoplasia, several studies indicate that levels of U34 tRNA modifications can be dynamically modulated^9,15,16^, which contradicts the notion that tRNA are constantly completely modified. For example, we showed that hormone signaling regulates the abundance of the catalytic subunit of the ELP complex, ELP3, thereby modulating U34 modification-levels in cancer cells^5^. Yet, the role of this activity in determining proteome composition in hormone-dependent cancers, and whether these cancers are addicted to high levels of U34-modified tRNA remains elusive. Nevertheless, these findings indicate that modulation of U34 tRNA modifications level may have a role in selectively regulating translation of specific transcript subsets, thereby dynamically reshaping the proteome in response to extracellular and intracellular cues. Mechanistically, such selective regulation is expected to be mediated via interactions between U34-modified tRNAs and cis-regulatory elements within target mRNAs, which ultimately determine their translational fate. Indeed, a recent study showed that levels of U34-modified tRNA can affect stability of mRNA harboring m□A modifications within codons requiring U34-modified tRNA for their decoding^17^. Additional studies have suggested that enrichment of U34-dependent codons within coding sequences^15,18^ and/or other encoded short peptides^19^ explain selective effects on proteome composition, but a unifying model of how tRNA modifications at the U34-position control proteome composition is lacking.

□

Therefore, here we systematically examined the link between alterations in U34-tRNA modifications, proteome composition and cellular phenotypes in prostate cancer cells. By applying our newly developed algorithm to model post-transcriptional regulation of gene expression^20^, we reveal a di-codon code that operates in concert with sequence contexts surrounding di-codons and features in the 5’ untranslated region (5’UTR) controlling translation initiation that collectively act as determinants for U34 modification-sensitive translation. Notably, genes with high occurrence of identified di-codons encode proteins associated with regulation of cell division across multiple species, which is consistent with the mitotic defects observed upon ELP3 suppression in prostate cancer cells. In contrast, non-transformed prostatic epithelial cells did not depend on U34-modified tRNA for their proliferation. Therefore, di-codons link U34 tRNA modifications to selective translation programs that facilitate cancer-associated phenotypes.

## RESULTS

### Prostate cancer cells require ELP3 to sustain proliferative potential

As discussed above, our previous studies suggested hormone-dependent regulation of gene expression via changes in tRNA modifications^5^. This raised the question whether hormone-dependent cancers are selectively sensitive to modulation of tRNA modifications. We therefore sought to determine whether modification of tRNA at the U34 position, mediated by the ELP3-ALKBH8-CTU1/2 axis, is required to sustain pro-cancer phenotypes in prostate cancer, and whether disruption of this axis selectively impairs cancer cells compared to non-transformed cells.

To this end, we reduced expression of ELP3 using CRISPR/Cas9 technology and two guide RNAs (gRNAs) in DU145 and LNCaP prostate cancer cell lines as well as the immortalized but non-transformed PNT1A prostatic cell line. While expression of ELP3 protein was higher in DU145 and LNCaP cancer cells as compared to PNT1A cells, all clones with ELP3-targeted gRNA showed reduced expression of the ELP3 protein as determined by western blotting (**Fig. 1A**). Expectedly, as quantified by mass spectrometry within the tRNA pool, this was associated with a decrease (but not elimination) of the mcm^5^s^2^U34 modification that requires activity of all enzymes in the ELP3-ALKBH8-CTU1/2 axis (**Fig. 1B**). The lack of complete elimination of the mcm^5^s^2^U34 modification in tRNA appears to be due to residual expression of ELP3 (**Fig. 1A** and see further below). Indeed, ELP3 has been shown to be an essential gene in mouse and complete abrogation of ELP3 may therefore not be achievable in some cell lines^21^. To determine whether reduced expression of ELP3 affected prostate cancer-associated phenotypes, we assessed ELP3-dependent proliferation. In contrast to non-transformed PNT1A cells, whose proliferation was insensitive to ELP3 suppression, both cancer cell lines showed a significant decrease in proliferation (**Fig. 1C**). Consistently, the colony formation assay in soft agar demonstrated a dramatic reduction in the number of colonies when ELP3 expression is suppressed in cancer cell lines (**Fig. 1D**); non-transformed PNT1A cells do not form colonies.

**Figure 1:**
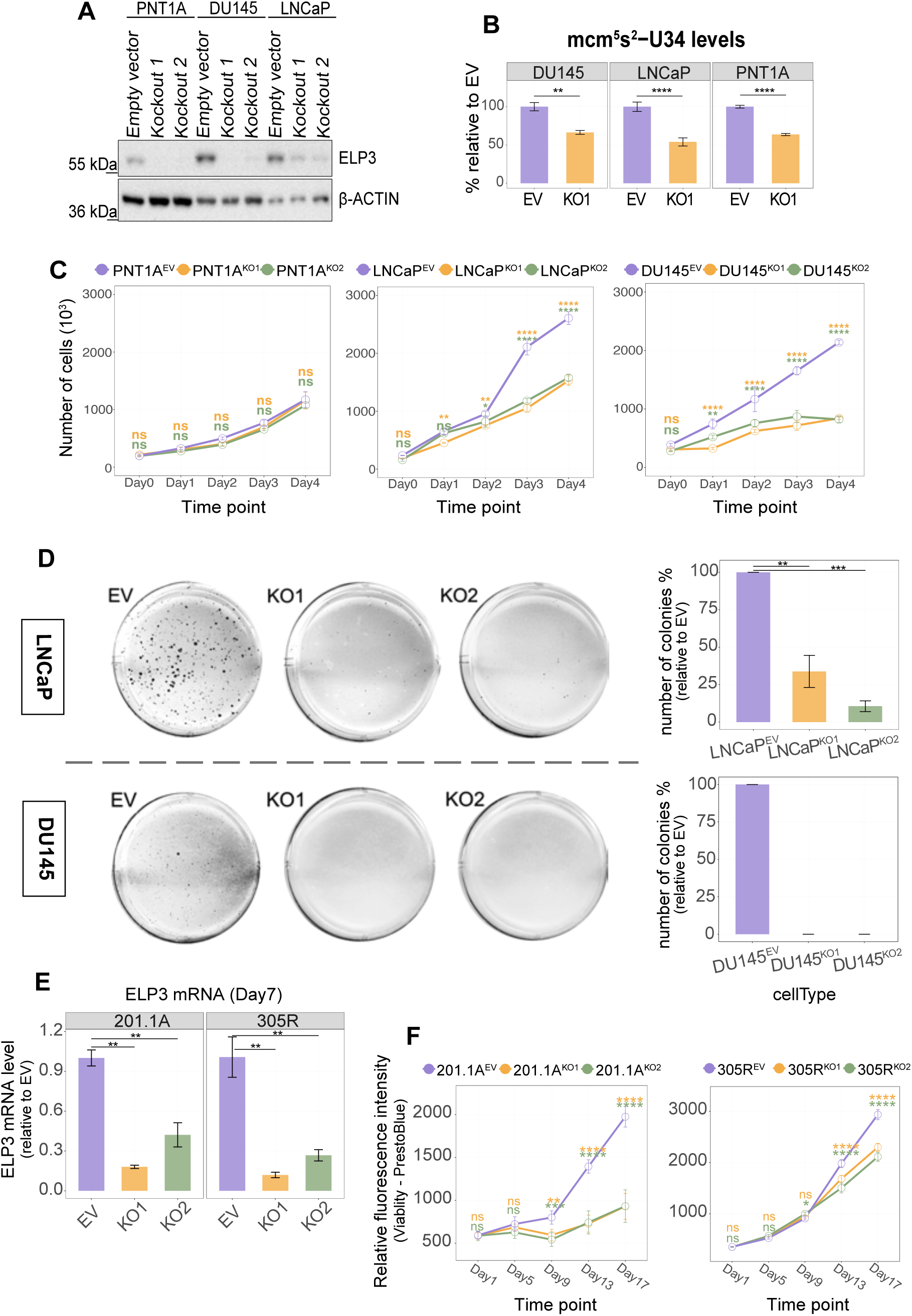
ELP3 is required to sustain proliferation of prostate cancer cell lines and organoids but not non-transformed cells. A) Western blot analysis for ELP3 in PNT1A, DU145 and LNCaP cells transduced with pLentiCRISPRv2 empty vector (EV), pLentiCRISPRv2 harboring ELP3 sgRNA #1 (Knockout1) and pLentiCRISPRv2 harboring ELP3 sgRNA #2 (Knockout2) (n = 3). β-actin was used as a loading control. B) Quantification of mcm^5^s^2^U levels, quantified using liquid chromatography–mass spectrometry (LC-MS), within the tRNA pool isolated from DU145^EV^ and DU145^KO1^ cells. Data represent mean ± SD of n=3 biological replicates. Statistical significance was assessed using Student’s t- test. **, p ≤ 0.01; ****, p ≤ 0.0001. C) Proliferation curves of PNT1A, LNCaP and DU145 cells transduced with empty vector (EV, purple), ELP3 sgRNA #1 (KO1; orange) and ELP3 sgRNA #2 (KO2; green). Each data point is mean ± SD of n=3 biological replicates. Statistical significance was assessed using two-way ANOVA. ns, not significant; *, p ≤ 0.05; **, p ≤ 0.01; ***, p ≤ 0.001; ****, p ≤ 0.0001. D) Representative images from soft agar colony formation assay of control and ELP3-depleted LNCaP and DU145 cells. Bar graphs show quantification of 3D colony numbers. Graphs represent mean ± SD of n=3 biological replicates. Statistical significance was assessed using a two-sided t-test. **, p ≤ 0.01; ***, p ≤ 0.001. E) RT-qPCR analysis of the 201.1A and 305R advanced prostate cancer organoids transduced with empty vector (EV, purple), ELP3 sgRNA #1 (KO1; orange) and ELP3 sgRNA #2 (KO2, green) 7 days post transduction. Data represent mean ± SD of 3 technical replicates. Statistical significance was determined using Student’s t-test (**, p ≤ 0.01). F) Organoid growth profiles of 201.1A and 305R lines transduced with empty vector (EV), ELP3 sgRNA #1 (KO1) and ELP3 sgRNA #2 (KO2). Each data point represents mean ± SD of n=5 biological replicates (2 technical replicates each). Statistical significance was assessed using two-way ANOVA. ns, not significant; *, p ≤ 0.05; **, p ≤ 0.01; ***, p ≤ 0.001; ****, p ≤ 0.0001.

To further determine the role of U34 modifications for prostate cancer proliferation, we used patient-derived organoid models of metastatic castration-resistant adenocarcinoma (201.1A) and neuroendocrine prostate cancer (305R)^22^. The same gRNAs and reagents used in prostate cancer cell lines (**Fig. 1A**) were employed to suppress expression of ELP3 in organoid models, resulting in reduced ELP3 mRNA levels as determined by qPCR (**Fig. 1E**). Consistent with findings in cell lines (**Fig. 1C**), organoid growth potential was sensitive to ELP3 suppression (**Fig. 1F**). Therefore, the proliferative capacity of prostate cancer cells depends on modification of tRNA at the U34 position mediated by the ELP3-ALKBH8-CTU1/2 axis.

### ELP3 is required for optimal elongation during translation of mRNAs encoding cell-cycle related proteins

To understand why prostate cancer cells depend on ELP3 for their proliferation, we studied ELP3-sensitive gene expression. Initially, we focused on the proteome and performed label-free quantitative proteomics on ELP3 suppressed DU145 (DU145^KO1/2^) and empty vector control (DU145^EV^) cell lines (i.e. from **Fig. 1A**; **Fig. 2A**) and quantified 4,351 proteins consistently across all cell lines and replicates. In agreement with residual activity of ELP3 in ELP3-suppressed cell lines (**Fig. 1A-B**), peptides from ELP3 were detected by mass spectrometry in all samples. Principal component analysis (PCA) revealed high reproducibility as samples clustered according to cell line in components one and two (**Fig. 2B**). However, DU145^KO2^ cells diverged from both DU145^EV^ and DU145^KO1^ cell lines in principal component 1 (**Fig. 2B**) in a fashion that could not be explained by a larger reduction in ELP3 expression in DU145^KO2^ relative to DU145^KO1^ cells, as DU145^KO1^ cells instead showed a more substantial reduction in ELP3 expression (**Fig. 2C**). In contrast, principal component 2 mirrored relative ELP3 expression between DU145^EV^ (most negative in component 2), DU145^KO2^ (intermediate in component 2) and DU145^KO1^ (most positive in component 2; **Fig. 2C**) cell lines. Therefore, although DU145^KO2^ cells are expected to reflect the ELP3-sensitive proteome, DU145^KO2^ cells may have acquired additional alterations to the proteome independent of ELP3 expression. We therefore focused our downstream analysis on DU145^KO1^ cells and used DU145^KO2^ cells to validate the findings.

**Figure 2:**
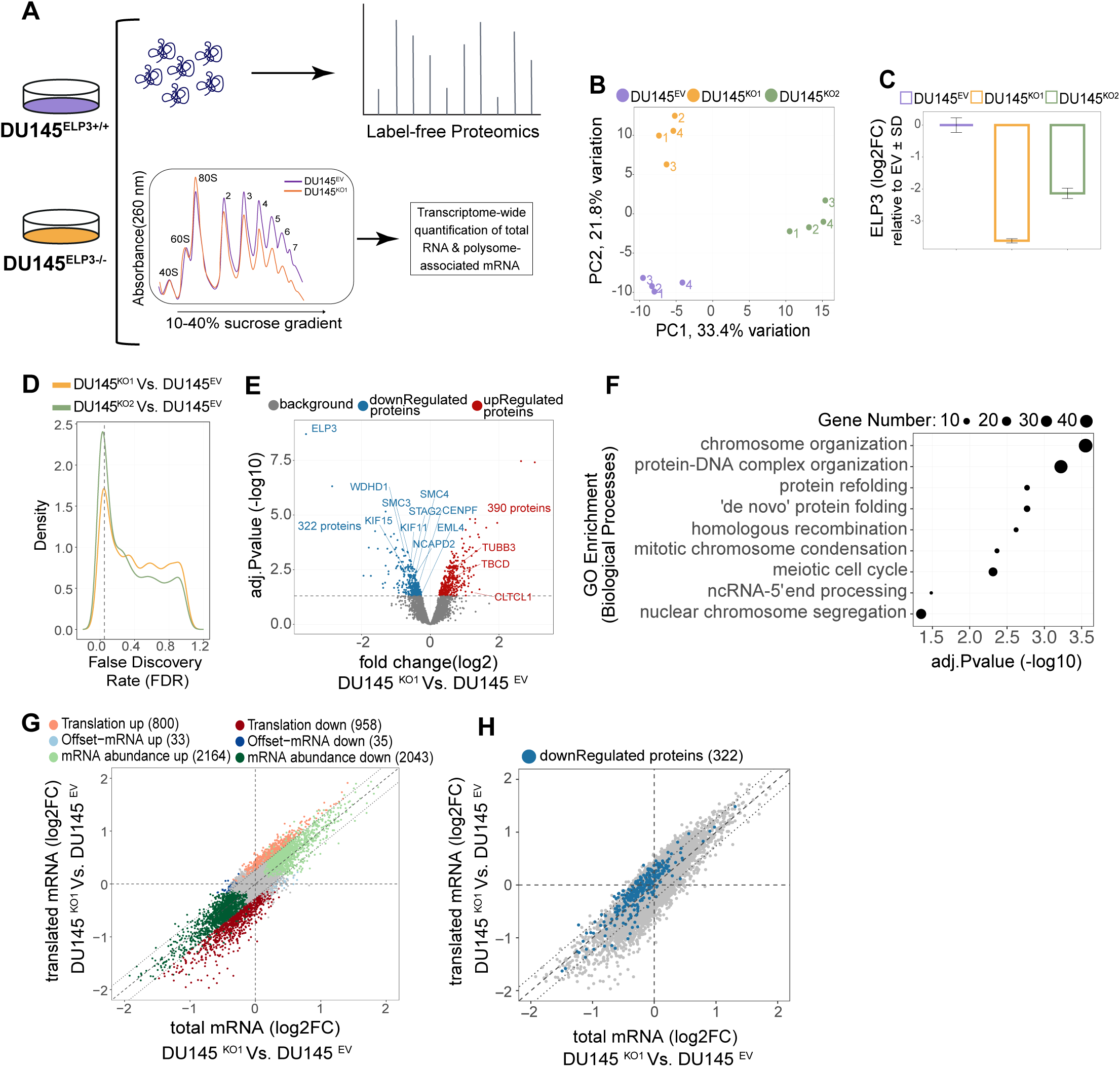
ELP3 is required for translation of mRNA encoding cell-cycle related proteins. A) Schematic of the experimental setup comparing empty vector control (DU145^ELP3+/+)^ to the ELP3-depleted cells (DU145^ELP3-/-^). Both cell lines were subjected to label-free proteomics and polysome profiling. For proteomics, global protein abundance was quantified by LC-MS/MS. During polysome-profiling, cytoplasmic extracts from indicated cell lines were fractionated on 10-40% sucrose gradients, and RNA was extracted from total lysates. Fractions were pooled to isolated polysome-associated mRNAs (mRNA associated with >3 ribosomes). Total and polysome-associated mRNA was quantified using RNA sequencing. B) Principal component analysis (PCA) of global proteome profiles in DU145^EV^, DU145^KO1^ and DU145^KO2^ cells. Each point represents one biological replicate. The first two principal components explain 33.4% and 21.8% of the total variance, respectively. C) ELP3 protein abundance measured by proteomics. Plotted are log₂ fold change relative to the empty vector control (mean ± SD across replicates). D) Kernel density distributions of false discovery rates (FDR) for differential protein abundance. A higher density of low FDR values reflects an increased number of significantly altered proteins upon ELP3 suppression. E) Volcano plot with log2 fold-changes on the x-axis compared to –log10 adjusted p-values on the y-axis to illustrate differential protein abundance in DU145^KO1^ relative to DU145^EV^. Each point represents one quantified protein, with significantly downregulated (blue, n=322) and upregulated (red, n=390) proteins highlighted (FDR < 0.05); non-significant proteins are shown in gray. F) Gene ontology (GO) enrichment analysis of proteins significantly decreased upon ELP3 suppression (DU145^KO1^ Vs. DU145^EV^). Enriched biological processes are shown, with significance determined after multiple testing correction (FDR < 0.05). G) Scatter plot comparing log₂ fold changes of polysome-associated versus total mRNA in DU145 cells (DU145^KO1^ Vs. DU145^EV^). Genes are classified by regulatory modes as determined by the anota2seq algorithm (FDR < 0.15), with colors indicating each category. The number of regulated transcripts per category is shown in the legend. H) Scatterplot as in Fig. 1G but highlighting mRNAs encoding proteins that were downregulated according to the proteomics analysis.

First, we used a Random Variance Model^23^ implemented in the anota2seq Bioconductor package^24^ to identify alterations in protein levels between DU145^KO1/2^ vs. DU145^EV^ cells. This revealed an enrichment of low p-value events (**Fig. 2D**) that allowed identification of hundreds of significantly (false discovery rate [FDR]< 0.05) up- or down-regulated proteins (**Fig. 2E** and **extended Fig. 2_01A**). In agreement with a common ELP3-sensitive proteome in DU145^KO1^ and DU145^KO2^ cell lines, proteins with altered expression in DU145^KO1^ cells were regulated in a similar fashion, albeit to a lesser extent, in DU145^KO2^ cells (**extended Fig. 2_01B**). This difference in magnitude is consistent with the relative reduction in ELP3 expression between DU145^KO1^ and DU145^KO2^ cell lines (**Fig. 2C**). Accordingly, suppression of ELP3 activity results in reproducible proteome alterations in prostate cancer cells.

We next sought to establish if the ELP3 sensitive proteome selectively targets cellular functions that may explain the observed reduction in proliferative capacity (**Fig. 1C-F**). To this end, we used Gene Ontology (GO) enrichment analysis for proteins that were up- or downregulated in the comparison of proteomes from DU145^KO1^ vs DU145^EV^ cells (i.e. from **Fig. 2E**). Notably, proteins whose levels decreased upon ELP3 suppression in DU145^KO1^ cells were enriched for cell cycle-related functions, in particular the mitotic process, and included *SMC2*, *STAG2*, *NCAPD2* and *NUSAP1* genes. Conversely, upregulated proteins were linked to basal cellular processes, including actin cytoskeleton organization (*CALD1*, *ARPC5* and *TPM1*) and ion homeostasis (*CALB2*, *ATP2B4*; **Fig. 2F**; **extended Fig. 2_01C**). Accordingly, the ELP3-sensitive proteome (**Fig. 2E-F**) is consistent with the observed reduced proliferative capacity in cells with suppressed ELP3 activity (**Fig. 1C-F**).

As ELP3 suppression resulted in reduced mcm^5^s^2^U modifications of tRNA, which is required for optimal decoding^5,8^ (**Fig. 1B**), it may alter protein levels by affecting translation elongation. A reduction in translation elongation is expected to lead to increased ribosome association with the mRNA in combination with reduced protein levels. To assess if reduced translation elongation underlies changes in proteome composition in ELP3-suppressed cells, we performed polysome profiling quantified with RNA sequencing in DU145^KO1^ and DU145^EV^ cell lines (**Fig. 2A**). During polysome profiling, the total mRNA pool is sedimented in a sucrose gradient depending on the number of bound ribosomes, thereby allowing isolation of mRNA associated with heavy polysomes (>3 ribosomes; hereafter referred to as polysome-associated mRNA)^25^. Total mRNA is isolated in parallel to determine if an alteration in polysome-associated mRNA represents a change in translational efficiency or upstream modulation of the total mRNA pool depending on e.g. altered transcription or mRNA stability. The obtained data set was of high quality as indicated by the sequencing depth and separation of samples according to RNA source (total or polysome-associated) and cell type in components one and two in a PCA analysis (**extended figure 2_01D-E).** In total, expression patterns of 10,448 genes were analyzed using the anota2seq algorithm, which identifies three modes of gene expression regulation: alterations in polysome-associated mRNA not explained by changes in mRNA levels (indicating a change in translational efficiency and referred to as “translation”); congruent changes in total and polysome-associated mRNA (referred to as “abundance”); and alterations in total mRNA not paralleled by corresponding changes in polysome-associated mRNA (referred to as “offsetting”). The latter gene expression mode is understudied but may reflect homeostatic processes^10^. Anota2seq analysis indicated that a larger proportion of genes showed a low p-value in the analysis of polysome-associated as compared to total mRNA changes in the comparison between DU145^KO1^ and DU145^EV^ cell lines (**extended figure 2_01F)**. Consistently, approximately 1,800 genes were classified as regulated via the translation mode by the anota2seq algorithm (**Fig. 2G**). In addition, a large subset of genes showed congruent alterations of polysome-associated and total mRNA, indicating ample changes in transcription and/or mRNA stability, while only a small fraction was translationally offset (**Fig. 2G**). To determine whether transcripts encoding proteins with reduced abundance upon ELP3-suppression also showed altered polysome-association, we assessed them in the anota2seq scatter plot (**Fig. 2H**). As evident by shift above the diagonal identity line in the anota2seq scatter, this showed that a majority of proteins with reduced abundance following ELP3 suppression have increased polysome-association relative to their total mRNA level, often in absence of total mRNA-changes, a pattern implying changes in elongation efficiency (**Fig. 2H**). Together, these data indicate that, in DU145 prostate cancer cells, ELP3 is required for optimal translation elongation of a subset of mRNAs encoding proteins involved in cell-cycle progression.

### Codon frequency poorly explains ELP3-sensitive protein expression

As the ELP3-ALKBH8-CTU1/2 axis modifies U34 in a subset of tRNA (U34-tRNA), most studies have assumed that the requirement for U34-tRNAs during decoding determines whether translation of an mRNA is sensitive to reduced levels of modified U34-tRNAs^5,16,19^. We therefore first assessed codon composition of mRNAs encoding proteins with reduced expression upon ELP3 suppression, which indicated that U34-codons (i.e. codons decoded by U34-tRNA) are overrepresented (including GAA, AAA, CAA, AGA and UUA) in this subset as compared to upregulated proteins in both DU145^KO1^ and DU145^KO2^ cell lines **(Fig. 3A; Extended Fig. 3_01A)**. Consistently, several indexes assessing codon bias, including codon adaptation index, differed between mRNAs encoding downregulated as compared to upregulated proteins **(Extended Fig. 3_01B)**. To account for a potential bias from amino acid composition, we normalized codon frequencies to amino acid counts and obtained similar results **(Extended Fig. 3_01C)**. Therefore, as expected, U34-codons are enriched among proteins downregulated upon ELP3 suppression in DU145 cells relative to their corresponding G-ending synonymous codons **(Extended Fig. 3_01D)**.

**Figure 3:**
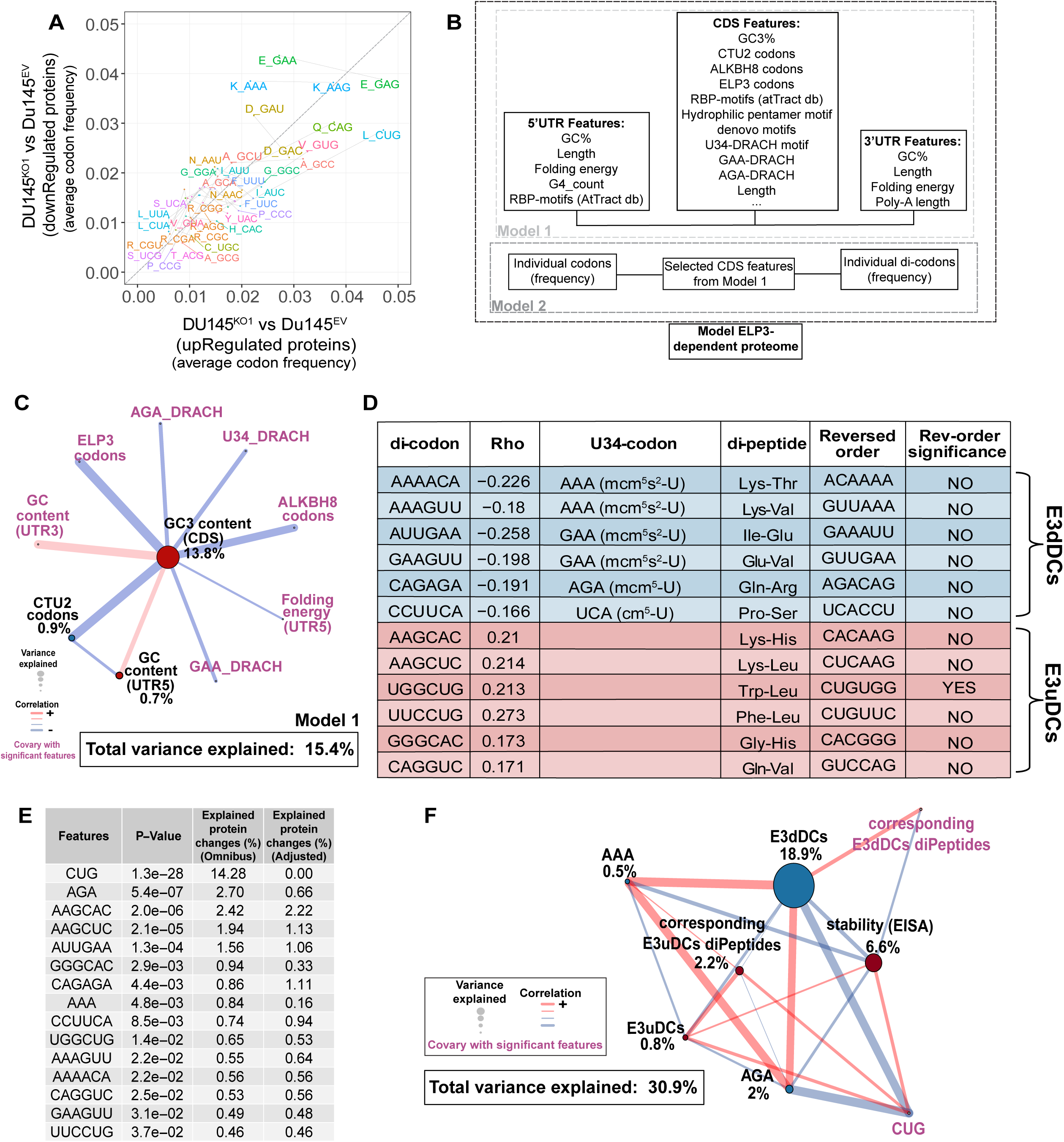
Di-codon, rather than single-codon, usage explains the ELP3-sensitive proteome. A) Comparison of codon frequencies between proteins down- and up-regulated upon ELP3 suppression in DU145 cells. Codons encoding the same amino acid are connected by gray lines and shown in matching colors. B) For modeling of the ELP3-sensitive proteome, cis-acting mRNA features were derived from 5′UTR, CDS, and 3′UTR sequences and integrated into Model 1 (FDR < 0.05). Model 2 combined the CDS features retained from Model 1 with individual codon and di-codon frequencies to model the ELP3-dependent proteome (FDR < 0.05). C) Network representation of Model 1 output. The total percentage of explained proteomic changes by the model is shown, together with additional breakdown by feature category. Features selected by the model are labeled in black while those in purple showed co-variance with selected features. Edges reflect the strength of the correlation between features, and node colors indicate association with down- or up-regulation following ELP3 suppression. D) Table summarizing di-codons identified in Model 2. Six di-codons associated with down-regulation were grouped as E3dDCs, while six associated with up-regulation were grouped as E3uDCs. Table columns for each di-codon report the correlation coefficient (r), whether a U34-decoded codon is present in the di-codon, the corresponding di-peptide, the reversed orientation of the di-codon together with a column indicating whether the reversed di-codon was significant as a factor explaining ELP3-sensitive protein expression in a univariate model (FDR < 0.05). E) Table summarizing features retained in Model 2. For each feature, the omnibus model p-value is shown, together with the proportion of regulation explained in the omnibus and adjusted model. The adjusted model indicates how much regulation each feature explains after adjustment for all other features. F) Extended model combining the features selected in Model 2 (FDR < 0.05) with EISA RNA stability scores and di-peptides derived from E3dDCs and E3uDCs. Node colors indicate associations with protein regulation, and edges represent correlations among features.

To determine how well codon composition explains the ELP3 sensitive proteome, we used our recently developed post-transcriptional network modelling method (postNet)^20^. Briefly, postNet assesses how a collection of features explains observed regulation patterns and determines the relationships between features. The output is a point estimate of the percentage of regulation explained by the whole model and each identified feature (”omnibus” output). Notably, as postNet applies a forward stepwise regression model, regulation explained by covariance between features will be assigned to the feature that can explain a larger proportion of the regulation. Therefore, postNet also assesses how much regulation each identified feature explains independently of all other identified features (”adjusted” output). The graphical output is a network with identified features as nodes and their correlations as edges. When applying postNet on alterations in protein levels between DU145^KO1^ and DU145^EV^ cells, we reasoned that elements within the untranslated regions (UTRs) may also influence how individual transcripts are regulated upon ELP3 suppression and therefore included features from both coding and UTR regions of the mRNA **(Fig. 3B; Model 1)**. For this analysis, to simplify interpretation, we created subsets of codons that are decoded by tRNA modified by only ELP3 (ELP3-codons), modified by both ELP3 and ALHBH8 (ALKBH8-codons), or modified by ELP3, ALKBH8 and CTU1/2 (CTU2-codons). Moreover, we assessed the hydrophilic pentamer amino acid motif reported previously to promote protein aggregation and degradation in the context of U34 codon-dependent translation elongation defects^19^. Although hydrophilic pentamer motifs were present in nearly 20% of the downregulated proteins and showed a small enrichment relative to non-regulated proteins **(Extended Fig. 3_01E)**, they did not reach statistical significance during postNet modeling (FDR > 0.05), suggesting that they do not play a major role in determining ELP3-sensitive gene expression. In total, approximately 15% of the ELP3-sensitive proteome was explained by a postNet model including GC3 content of the CDS (i.e. the GC content in the 3rd position of the codon that correlates with the presence of U34-codons), the frequency of CTU2-codons, and the GC content of the 5’UTR **(Fig. 3C; Extended Fig. 3_01F)**. This suggests that both 5’UTR and coding regions include features that independently underlie the ELP3-sensitive proteome, possibly via an interplay between translation initiation and elongation mechanisms. Although a postNet model is not expected to fully explain observed regulation due to e.g. measurement error and indirect effects, 15% was lower than what we observed in our previous studies^20^. Accordingly, the limited ability of codon frequencies in explaining the ELP3-sensitive proteome suggests the existence of additional mechanisms controlling ELP3-sensitive protein expression.

### Altered mRNA stability independently contributes to the ELP3-sensitive proteome

To further explore features affecting ELP3-sensitive protein synthesis, we considered a recently published mechanism for mcm^5^s^2^U34 dependent regulation of gene expression^17^. Codons carrying m□A marks are inefficiently decoded by the translation machinery, rendering them suboptimal and prone to inducing ribosome collisions^26^, which can trigger downstream mRNA decay. The mcm□s²U34 modification in the tRNA anticodon loop was shown to oppose this by facilitating translation of m□A-modified U34-codons and thereby stabilizing the mRNA. To determine whether this mechanism contributes to the observed regulation, we added U34-decoded DRACH motifs (which are potential sites for m□A modifications) along the CDS to the model **(Fig. 3B)**. Although these motifs were significantly enriched among downregulated proteins **(Extended Fig. 3_01G)**, which is expected given the observed codon bias **(Fig. 3A)**, they did not independently explain ELP3-sensitive protein expression **(Fig. 3C)**. Nevertheless, to further assess whether reduced protein levels upon ELP3-suppression were driven by altered mRNA decay rather than impaired protein synthesis alone, we applied Exon-Intron Split Analysis (EISA)^27^. EISA distinguishes transcriptional from post-transcriptional regulation of mRNA levels by comparing the abundance of exonic and intronic reads from RNA sequencing data. Intronic reads are assumed to primarily reflect nascent, unspliced transcripts and are therefore considered as a proxy for transcriptional activity, while exonic reads represent the mature, processed transcript. In this framework, a stronger reduction in exonic signal relative to intronic signal is interpreted as selective mRNA destabilization **(Extended Fig. 3_02A)**. Notably, 10% of the proteins downregulated upon ELP3 suppression (32 out of 322) corresponded to transcripts classified as significantly unstable upon ELP3-suppression by the EISA method **(Extended Fig. 3_02B)**. Furthermore, an empirical cumulative distribution function (ECDF) analysis revealed a subtle yet significant shift in protein fold-change values towards reduced protein levels for EISA-identified destabilized transcripts compared to the background **(Extended Fig. 3_02C)**. Importantly, there is a negative correlation between mRNA stability and the frequency of U34 tRNA-decoding DRACH motifs along the CDS (Pearson’s ρ = –0.23, pValue = 5.44 × 10⁻□□), indicating that a higher number of motifs is associated with reduced mRNA stability **(Extended Fig. 3_02D)**. Furthermore, there was a trend for a global reduction in m□A among poly-adenylated mRNA in DU145^KO1^ and DU145^KO2^ as compared to DU145^EV^ cells **(Extended Fig. 3_02E)**. These observations are consistent with the reported link between m□A and mcm□s²U34 modifications in translation dynamics^17^ and suggests that the impact from this mechanism may have been more substantial if m□A levels had been maintained upon ELP3-suppression. Incorporating the EISA stability metric into postNet modeling increased the percentage of explained regulation from ∼16 to ∼23% **(Extended Fig. 3_02F)** and eliminated the contribution of CTU2-codon frequency from the model, which is consistent with the correlation between U34-DRACH frequency and EISA estimated stability. Importantly, the variance explained by mRNA stability remained unchanged after adjusting for the contributions of GC3 content of the CDS and the GC content of the 5’UTR **(Extended Fig. 3_02F)**. Therefore, altered stability of the mRNA seems to be a distinct and independent mechanism for controlling ELP3-sensitive protein synthesis.

### Di-codon code and context explain ELP3-sensitive translation

Given that GC3 content explained the largest proportion of regulation in the postNet model also incorporating mRNA stability **(Extended Fig. 3_02F)**, we considered that there could also be overlooked features in the coding region determining ELP3-sensitive protein expression. We therefore assessed whether adjacent codons (di-codons), presumably positioning tRNA in the P-site and A-site and thereby influencing translation elongation, may underlie ELP3-sensitive translation^28^. Surprisingly, postNet analysis identified 12 di-codons as well as three codons that together explained approximately 30% of ELP3-sensitive protein expression **(Extended Fig. 3_03A)**. Six dilZcodons were enriched among mRNAs encoding proteins with reduced level upon ELP3-suppression **(Fig. 3D**, referred to as ELP3 down di-codons “E3dDCs”; **Extended Fig. 3_03A** [blue circle]; **Extended Fig. 3_03B)**, each containing a U34-codon. An additional 6 di-codons were enriched among mRNAs encoding up-regulated proteins **(Fig. 3D**, “E3uDCs”**; Extended Fig. 3_03A** [red circle]; **Extended Fig. 3_03B)**. Importantly, E3dDCs and E3uDCs were also enriched in the ELP3-sensitive proteome obtained from DU145^KO2^ cells **(Extended Fig. 3_03C)**, albeit with reduced magnitude, consistent with the partial (∼50%) suppression of ELP3 in DU145^KO2^ cells **(Fig. 2C)**. Strikingly, after adjusting each postNet-identified codon or di-codon (i.e. adjusted model), the regulation explained by individual codons was almost completely lost, which was in contrast to di-codons **(Fig. 3E)**. This implies that di-codons provide unique information not captured by individual codons and that individual codons co-vary with occurrence of di-codons. Furthermore, incorporating the frequency of di-peptides encoded by the selected di-codons only marginally improved the proportion of explained regulation (∼1%). In support for that the order of the codons within E3dDCs/E3uDCs is relevant, only one di-codon was significant in univariate analysis within postNet after reversing the order **(Fig. 3D),** despite having comparable frequencies across all CDSs **(Extended Fig. 3_03D)**. To assess whether E3dDC frequency influences mRNA stability, we calculated their correlation for the set of proteins detected in the proteomics experiment. This indicated a substantially lower correlation between E3dDCs and EISA stability than that observed between EISA stability and the U34-DRACH motif **(compare Extended Fig. 3_02D and Extended Fig. 3_03E)**. Consistently, E3dDC frequency remained the feature explaining the largest percentage of ELP3-sensitive protein expression also after including EISA stability **(Fig. 3F; Extended Fig. 3_03F)**. In aggregate, these findings support a model in which specific di-codon configurations contribute to ELP3-dependent regulation of protein synthesis by shaping elongation dynamics and that this occurs independently of, but in parallel with, mRNA stability-dependent modulation of the proteome.

### E3dDC-associated failure in activating the c-ISR translation program during ELP3 suppression

The canonical Integrated Stress Response (c-ISR) modulates both global and selective mRNA translation in response to various cellular stresses^29^. Stresses include perturbations in the translation elongation process that cause ribosome stalling and collisions, which can be recognized by stress-sensing kinase GCN2. During c-ISR, stress sensing kinases phosphorylate eIF2α at Ser51, leading to a global reduction in protein synthesis and selective translation of mRNAs harboring e.g. upstream open reading frames (uORFs), including ATF4. Consistent with activation of this pathway upon ELP3-suppression, western blot analysis revealed increased phosphorylation of eIF2α as well as increased expression of ATF4 in DU145^KO1^ and DU145^KO2^ cells **(Fig. 4A)**.

**Figure 4:**
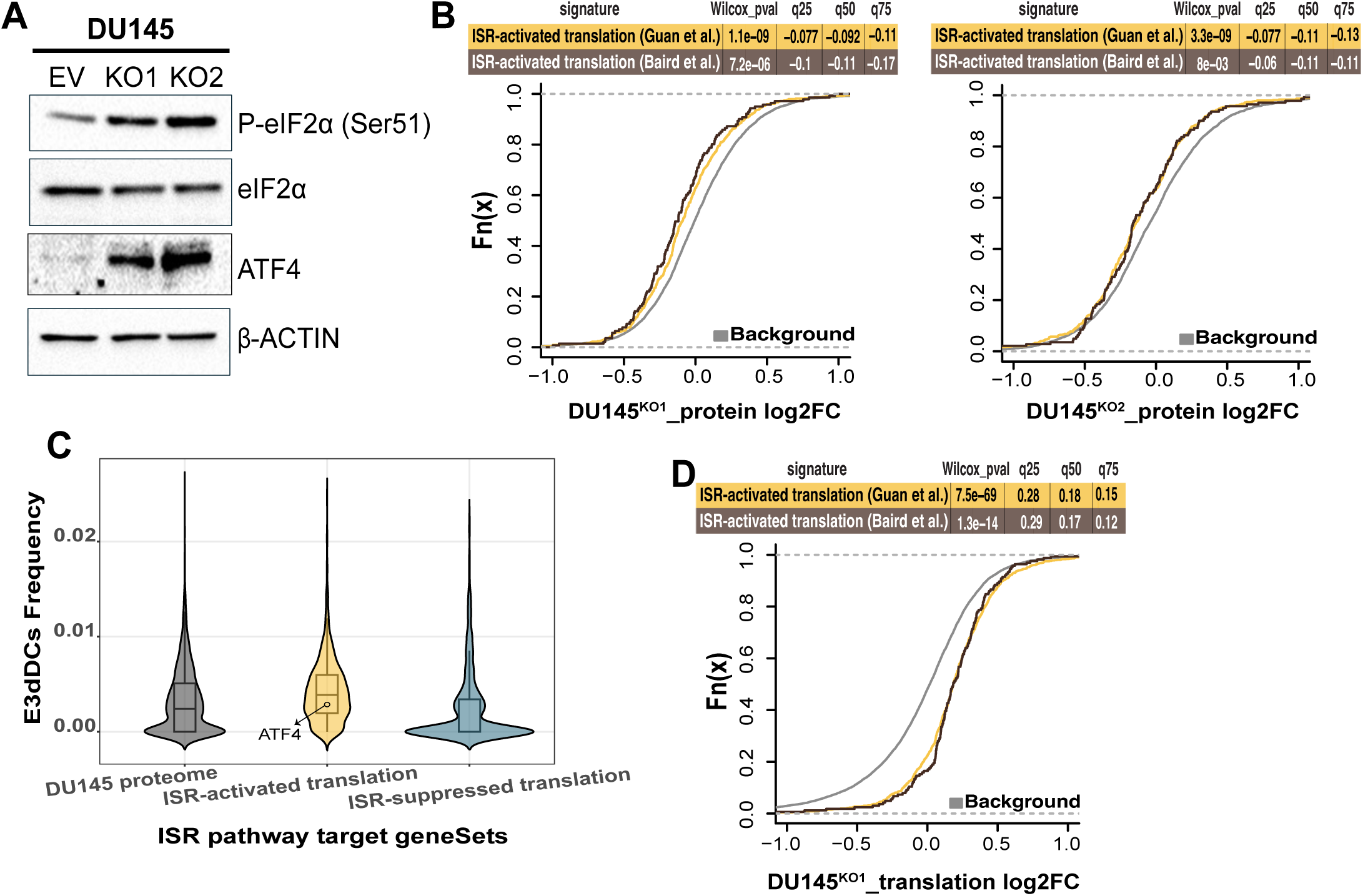
E3dDC-associated failure in activating the c-ISR translation program during ELP3 suppression. A) Western blot analysis of phosphorylated eIF2α (Ser51), total eIF2α and ATF4 were assessed in Empty vector (EV), DU145^KO1^, and DU145^KO2^ cells, with β-actin used as a loading control (n=3). B) Empirical cumulative distribution functions (ECDFs) of protein abundance changes comparing two independent ISR-activated signatures in DU145^KO1^ (left) and DU145^KO2^ (right) to the background. Each signature collects transcripts reported as translationally upregulated upon ISR activation. Statistical significance was assessed by two-sided Wilcoxon rank-sum test, with p-values and distribution shifts at the 25th, 50th, and 75th percentiles indicated. C) Violin plots of E3dDC frequencies across the DU145 proteome compared to ISR-signatures with genes reported as translationally activated or suppressed. Boxplots indicate medians and interquartile ranges. ATF4, a canonical ISR target, is marked. D) Same analysis as in (B), but showing empirical cumulative distribution functions (ECDFs) of log₂ fold-changes in polysome-association adjusted for total mRNA (“Translation”) from anota2seq analysis of DU145^KO1^ versus DU145^EV^.

However, in contrast to the expected increased expression of genes whose translation is activated during c-ISR (“c-ISR signature”^30,31^), we observed a marginally reduced protein expression of this gene set in both DU145^KO1^ and DU145^KO2^ cells **(Fig. 4B)**. A closer examination revealed that genes within the c-ISR signature are enriched for E3dDCs relative to all proteins detected in DU145 cells and transcripts whose translation is suppressed during c-ISR **(Fig. 4C)**. Consistent with a key role in E3dDCs in determining expression of c-ISR signature-genes, binning of genes within the c-ISR signature according to their E3dDC frequency revealed that reduced protein expression paralleled E3dDC frequency **(Extended Fig. 4_01A)**. Moreover, the increased expression of ATF4 **(Fig. 4A)** could be explained by the relatively lower frequency of E3dDCs as compared to other genes in the c-ISR signature **(Fig. 4C)**. In agreement with translation elongation-dependent modulation of ISR-signature expression, these genes showed increased polysome-association **(Fig. 4D)** despite the marginal reduction in protein level **(Fig. 4B)**, and binning c-ISR signature-genes based on E3dDC frequency revealed that mRNAs with higher E3dDC frequency were associated with polysomes to a larger extent **(Extended Fig. 4_01B)**.

In summary, despite activation of c-ISR, genes within the c-ISR signature that would be expected to show translational activation showed paradoxically reduced protein expression upon ELP3 suppression. This appears to be a consequence of high E3dDC frequency among c-ISR signature genes affecting translation elongation. Therefore, E3dDC context overrides activation of c-ISR-associated translation programs.

### E3dDCs have a distinct distribution between genes and within the CDS

As E3dDCs are completely absent from a large proportion of all genes **(Extended Fig. 3_03D)** and show higher frequencies among genes from the c-ISR signature **(Fig. 4C),** their distribution appears to be non-random across genes. To better understand the underlying properties of E3dDCs, we first sought to determine whether the overall frequency of E3dDCs is higher or lower than would be expected by the frequencies of the individual codons. To this end, we calculated the Poisson deviation between observed and expected E3dDC frequencies predicted by a zero-order Markov model across all genes using consensus coding sequences (CCDS; **Fig. 5A)**. This showed that, despite being absent from a large proportion of mRNAs, E3dDC consistently ranked within the top deciles for Poisson deviation. Importantly, this is not observed for the corresponding dipeptides encoded by E3dDCs **(Extended Fig. 5_01A)**. The higher frequency of E3dDCs is consistent with E3dDCs being selectively incorporated for their ability to regulate translation. Further supporting a role in gene expression regulation, the number of E3dDCs (assessed via bins) does not parallel the length of the coding region **(Extended Fig. 5_01B)**, which is contrast to U34-codons. As E3dDC frequency does not parallel length of CDS, but instead has a role in regulating gene expression, we considered that they may be enriched in mRNAs encoding proteins with specific functions. To explore this, we analyzed the functional enrichment among proteins encoded by transcripts containing multiple E3dDCs in human, mouse, and yeast. This revealed a substantial overlap in enriched categories among genes carrying multiple E3dDCs between species **(Fig. 5B)**. In all species, the top-enriched terms were consistently associated with cell cycle regulation and mitotic processes. In contrast, transcripts lacking E3dDCs were for example consistently enriched in immune and metabolic processes **(Extended Fig. 5_01C)**, indicating a functional segregation in E3dDC usage that may reflect distinct modes of translational regulation among gene classes.

**Figure 5:**
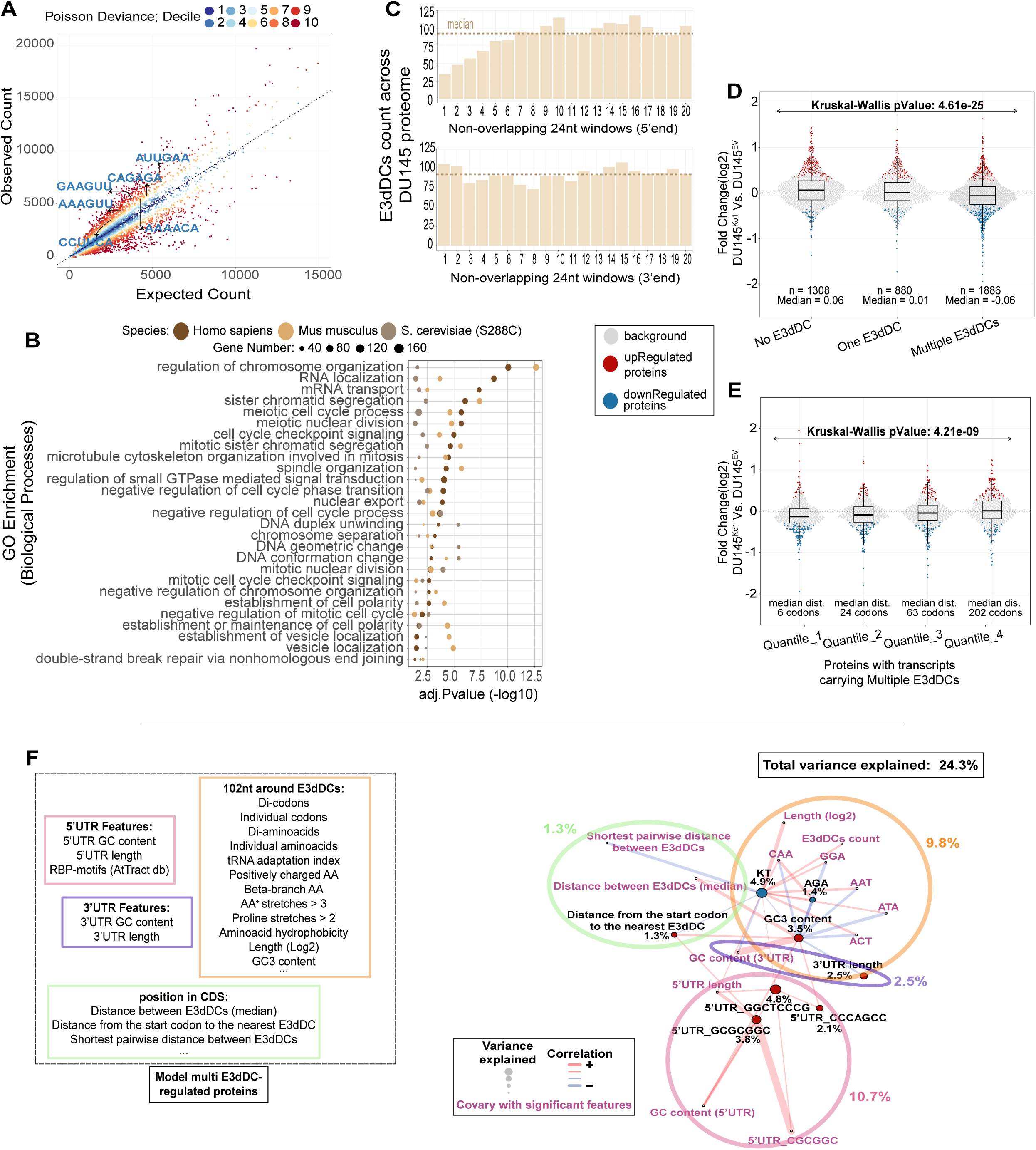
E3dDC context affects sensitivity to ELP3-dependent translation defects. A) Scatterplot of observed versus expected di-codon counts from the longest CDS isoforms in the CCDS database. Expected counts were derived from a zero-order Markov model, in which di-codon frequencies are predicted as the product of the independent single-codon frequencies, assuming no positional dependence between adjacent codons. Each point represents one di-codon, colored by Poisson deviance decile (dark blue = low, dark red = high). The diagonal line indicates equality between observed and expected counts. Labeled points correspond to E3dDCs. B) Cross-species GO enrichment analysis of transcripts carrying multiple E3dDCs in their CDS. Shown are GO biological processes (BP ontology) that were significantly enriched in all three species (FDR<0.05). Dot size indicates the number of genes, and color denotes species origin. The x-axis represents enrichment significance (–log10 adjusted p-value). Enriched GO terms are ranked according to adjusted significance values obtained from the human dataset. C) Meta-analysis of distribution of E3dDCs across detected DU145 proteins. Coding sequences were divided into consecutive non-overlapping 24-nt windows from the 5′ (top) and 3′ (bottom) ends. Bars represent the total number of E3dDCs across all DU145 proteins within each window, and the dashed line marks the median count across windows. D) Beeswarm plots of protein log₂ fold-changes (DU145^KO1^ Vs. DU145^EV^), within the entire DU145-detected proteome grouped by E3dDC content in the CDS: no, a single or multiple E3dDCs. Upregulated (red) and downregulated (blue) proteins are highlighted, and non-regulated proteins (FDR > 0.05, ANCOVA) are shown in gray. Boxplots indicate distribution and median values, with group sizes (n) and medians annotated. Statistical differences across groups were assessed using the Kruskal–Wallis test (p-value = 4.61e-25). E) Same analysis as in (D), but restricted to proteins with more than one E3dDC in their CDS. For each transcript, pairwise distances between E3dDCs were calculated, and the minimum distance was used. Proteins were then stratified into quartiles based on the minimum distance between E3dDCs. Within each quartile, upregulated and downregulated proteins are shown separately, while non-regulated proteins (FDR > 0.05, ANCOVA) are indicated in gray. Median distances between E3dDCs are indicated. Statistical differences across groups were evaluated using the Kruskal–Wallis test (p-value = 4.21e-09). F) (Left) Overview of cis-acting features investigated in the local vicinity of E3dDCs to model regulation of multi-E3dDC proteins. Features were derived from 102nt windows surrounding E3dDCs within a CDS. Categories include sequence composition (e.g., codons, amino acids, di-peptides), biochemical properties (e.g., hydrophobicity), and E3dDCs positional metrics (e.g., distance of the first E3dDC from the start or inter-E3dDC spacing). Features such as di-codons, individual codons, di-amino acids, individual amino acids, and RBP motifs (ATtRACT database) were pre-filtered, retaining only those significant at FDR < 0.15. The final model was then constructed using the same FDR threshold. (Right) Network representation of local feature modeling in multi-E3dDCs protein group. The proportion of proteome regulation explained by the model is shown, with contributions from different regions of the mRNA indicated. Features labeled in black collectively were selected by the modeling while those in purple show co-variance. Edges represent correlations between features, while node colors indicate associations with down- or up-regulation of proteins with multiple E3dDCs following ELP3 loss.

To determine whether the spatial distribution of E3dDCs within the coding region follows any pattern, we partitioned the CDS of all proteomics-detected proteins into consecutive, non-overlapping 24 nucleotide bins, starting from either the 5’ or 3’ end of the CDS, and pooled E3dDC frequency within bins across proteins. This identified a depletion of E3dDCs near the 5′ end of the CDS followed by a largely uniform distribution across the remaining CDS **(Fig. 5C)**. Interestingly, a similar pattern was observed in mouse but not in *S. cerevisiae* **(Extended Fig. 5_01D)**. The underrepresentation of E3dDCs in the beginning of the CDS may serve to prevent elongation stalling in the beginning of the coding sequence^32^.

In aggregate, E3dDCs appear at a higher frequency than expected, their frequency does not mirror the length of the CDS but parallels activity of encoded proteins in biological processes, and they are depleted from the translation ramp, all consistent with a role in regulating mRNA translation.

### E3dDC context underlies ELP3-dependent translational defects

As indicated above, for the ISR signature, the number of E3dDCs appears to affect the extent of reduction in the corresponding protein **(Extended Fig. 4_01A)**. To systematically assess this, we binned transcripts by the number of E3dDCs they contained and examined the associated protein fold changes. This analysis revealed an inverse relationship, whereby mRNAs harboring more than one E3dDC showed greater reduction in protein abundance compared with those lacking E3dDCs **(Fig. 5D)**. These findings indicate a frequency-dependent impact of E3dDCs on translational efficiency. The higher frequency of E3dDCs being associated with reduced protein expression upon ELP3-suppression may depend on their relative position within the CDS. To assess this, we investigated whether the distance between E3dDCs affected sensitivity to ELP3 suppression. To this end, we calculated all pairwise distances between E3dDCs within each CDS and related the smallest distance with protein fold changes. Interestingly, transcripts where E3dDCs are positioned closer to each other showed larger reductions in protein abundance upon ELP3 suppression **(Fig. 5E)**. These results indicate that the E3dDC context plays a critical role in determining susceptibility to ELP3-dependent translational defects. To explore this further, we focused on the subgroup of proteins with multiple E3dDCs that still exhibited a range of fold-change differences in protein expression upon ELP3 suppression **(Fig. 5D)** and used postNet together with a large set of features surrounding the E3dDCs as well as features from the UTRs as input **(Fig. 5F left)**. The resulting postNet model could explain about 24% of the variation in expression for the subset with multiple E3dDCs and notably indicated features from the 5’UTR as the major factor determining ELP3-sensitivity together with e.g. the distance between the most upstream E3dDC and the start of the coding sequence **(Fig. 5F right; extended Fig. 5_02A and B)**.

The identification of the 5’UTR as a key feature determining the impact of E3dDCs **(Fig. 5F)** suggests that higher or lower initiation of translation could affect whether an E3dDC will lead to translation defects or not. In support of this model, residuals from a linear regression between polysome association and total mRNA (adjusted for CDS length) in control cells (EV) showed a significant negative correlation with E3dDCs frequency **(Extended Fig. 5_01F)**. This indicates that transcripts with multi-E3dDCs recruit ribosomes less efficiently than expected under control conditions, pointing towards coordination between initiation and elongation steps. Notably, human and mouse transcripts carrying multiple E3dDCs exhibit significantly longer 5′UTRs compared to transcripts lacking E3dDCs **(Extended Fig. 5_01G)**. As 5′UTR length is a major determinant of translation initiation also underlying dynamic modulation of translation^33^, it suggests a context-dependent interaction between translation initiation and E3dDC-dependent translation elongation defects.

Therefore, the context of E3dDCs, including features in the 5’UTR affecting translation initiation as well as CDS features surrounding E3dDCs, tune how sensitive mRNAs with multiple E3dDCs are to ELP3-supression.

### E3dDC regions modulate translation elongation depending on sequence context and ELP3-activity

Our findings from above suggest that the context of E3dDCs modulate their ability to regulate translation elongation downstream of the activity of the ELP3-ALKBH8-CTU1/2 pathway. To evaluate this, we focused on the coding regions of STAG2 and TRMT6 that both encode proteins downregulated following ELP3 suppression but with different mechanisms according to polysome-profiling. Specifically, while reduced expression of STAG2 protein was associated with increased polysome-association, which is in agreement with inhibition of translation elongation, TRMT6 showed a congruent reduction in mRNA level and polysome-association, consistent with reduced mRNA abundance **(Fig. 6A)**. Moreover, although both genes contain multiple E3dDCs, the contexts are different as STAG2 has multiple E3dDCs in close proximity **(Fig. 6B)**, which was identified as a context associated with higher sensitivity to ELP3-supression **(Fig. 5E)**. To evaluate whether E3dDC sequence contexts mediate these differences in regulation, we selected the region containing four E3dDCs as well as an upstream length-matched control region from STAG2, and a region containing one E3dDC together with a length-matched region in between the two E3dDCs from TRMT6 **(Fig. 6B)**. Notably, the selected E3dDC-region of TRMT6 has a slightly higher frequency of U34-codons than the corresponding region of STAG2 while the full length CDS regions have comparable U34-codon frequencies **(Fig. 6C)**. The selected regions were cloned into a P2A-flanked monocistronic dual-fluorescence reporter in between an upstream GFP and a downstream mCherry. Accordingly, a reduction in mCherry after normalization to GFP is consistent with reduced translation elongation of the inserted sequence. As controls, we generated reporters with an efficiently translated “linker” sequence or a poly-lysine (“polyK”) sequence that leads to defects in translation elongation^34^. Expression of reporters was analyzed by flow cytometry focusing on cells with mid-to-high and high expression **(Extended Fig. 6_01A)**.

**Figure 6:**
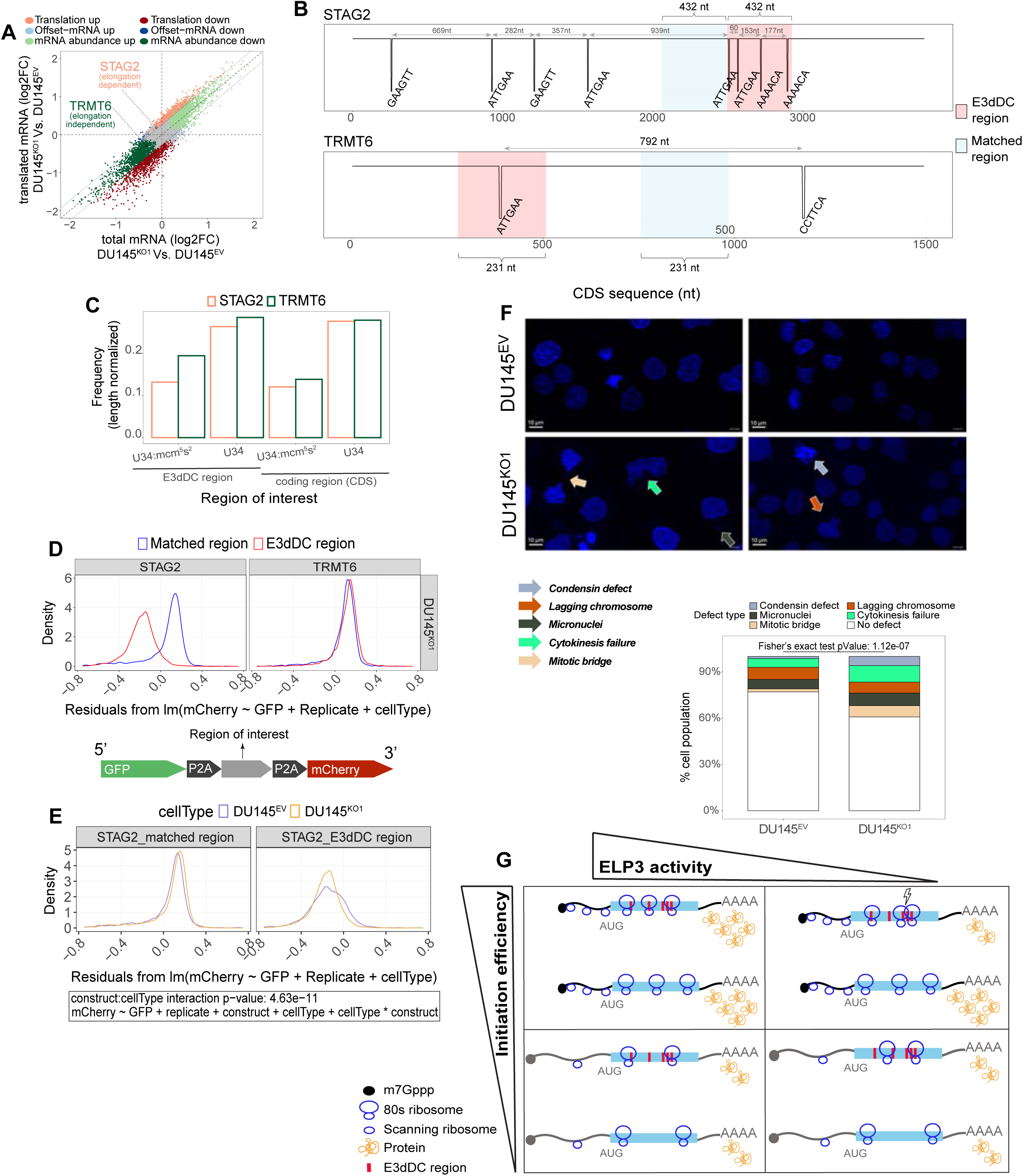
E3dDC regions act as translation traps, sensing levels of U34-modified tRNA in a context dependent fashion. A) Same as Fig. 2G but with TRMT6 and STAG2 highlighted as examples–of proteins encoded by transcripts containing multiple di-codons. Both proteins are significantly downregulated upon ELP3 suppression, but via distinct regulatory modes: TRMT6 falls into the mRNA abundance mode, showing concordant decreases in total and ribosome-associated mRNA, whereas STAG2 represents the translation mode, characterized by altered ribosome association without corresponding changes in mRNA abundance. B) Coding sequences of STAG2 and TRMT6 indicating E3dDCs. E3dDC-regions (red) and sized-matched control regions lacking E3dDCs (blue), were inserted into dual-fluorescence reporter constructs. C) Frequency of U34 ^cm5/mcm5/mcm5s2^ and U34^mcm5s2^ -decoded codons in STAG2 and TRMT6. Bars show length-normalized counts within selected E3dDC-regions and across the CDS. D) Density plots of per-cell–residuals (from model: mCherry ∼ GFP + replicate + cell type) for STAG2 and TRMT6 in DU145^KO1^ cells (a higher residual corresponds to higher expression of mCherry after normalizing to GFP-expression). Residual distributions from cells expressing reporters with E3dDC-regions (red) are compared to those from cells expressing–control-regions (blue).–(bottom) Overview of the sequence design for the dual-fluorescent reporter. E) Similar as (D) but for STAG2-regions in DU145^EV^ vs DU145^KO1^ cell lines. Residuals are shown separately for DU145^EV^ and DU145^KO1^ cells expressing the control-region (left) and the E3dDC-region (right). A significant construct:cell type interaction (p = 4.63e–11) indicates that the E3dDC- and control-regions differ in expression depending on cell type. F) Live-cell imaging showing representative frames of DU145^EV^ and DU145^KO1^ cells undergoing mitosis. Mitotic defects are indicated by arrows of different colors. Data for mitotic defects were collected from 10 imaged fields from DU145^EV^ and 13 imaged fields from DU145^KO1^ cells during–time-lapse recordings (8-min intervals over 4.8 h) across multiple wells. Mitotic defects were summarized in a bar graph and Fisher’s exact test was used to assess a difference in the proportion of mitotic defects depending on cell type. G) Model for how E3dDC regions control translation elongation dynamics depending on ELP3-activity, affecting levels of U34-modified tRNA, and efficiency of translation initiation. Limited amounts of U34-modified tRNA causes translation defects under conditions when initiation rate is high, which in turn leads to reduced levels of encoded proteins.

Flow cytometry analysis showed that in DU145^KO1^ cells, the GFP-normalized mCherry signal from the reporter containing the STAG2 E3dDC region was markedly reduced compared to the matched region **(Fig. 6D)**, indicating that translation elongation was impaired by the STAG2 E3dDC-region. This was in contrast to TRMT6, where the normalized mCherry signals from the TRMT6 E3dDC and matched regions were similar (**Fig. 6D**). To determine whether E3dDC regions act as sensors for ELP3-activity, we compared expression of STAG2 and TRMT6 E3dDC reporters depending on ELP3-status by evaluating reporter expression in both DU145^KO1^ and DU145^EV^ cells. This revealed a reduction of normalized mCherry signal in DU145^KO1^ relative to DU145^EV^ cells for the STAG2, but not the TRMT6, E3dDC reporter (p-value = 4.63e-11; **Fig. 6E**). In aggregate, these findings indicate that difficult-to-translate E3dDC containing-regions modulate translation elongation downstream of the ELP3-ALKBH8-CTU1/2 axis.

### Mitotic defects in DU145^KO1^ cells upon suppression of ELP3 activity

As indicated above, E3dDCs are enriched in mRNAs encoding cell cycle regulators, in particular those affecting mitosis. To evaluate if this could underlie the reduced proliferative capacity observed in prostate cancer cells **(Fig. 1C)**, we performed live cell imaging of DU145^KO1^ and DU145^EV^ **(Fig. 6F; Extended Fig. 6_01B)**. This identified that mitosis defects were markedly more frequent in DU145^KO1^ cells (p = 1.12e-07) and included uneven distribution of chromosomes, prolonged mitotic bridge and poor chromosome aggregation leading to presence of lagging chromosomes and micronuclei, as well as condensin defects **(Fig. 6F)**. These findings support that ELP3 regulates translation elongation of E3dDC-enriched mRNAs encoding proteins involved in mitosis, thereby orchestrating the cell division process.

## DISCUSSION

tRNAs have traditionally been assumed to be fully modified without dynamic regulation to ensure optimal decoding. In contrast, several recent studies have indicated elevated modifications of tRNA at the U34 position in cancer^15,16,35^ and we previously showed that the level of U34-modified tRNA is tuned by steroid hormone signaling in cancer cells^5^. However, the role of such modifications in establishing the cancer cell proteome, and whether hormone-driven cancers depend on this mechanism to maintain cancer-associated phenotypes was unclear. This study documents a key role of U34-modifications in sustaining proliferative potential in prostate cancer, but not in non-transformed, cells (**Fig. 1**). Although additional studies are required, these findings point towards targeting the ELP3-ALKBH8-CTU1/2 axis as a potential approach for treatment of hormone-driven cancers.

Post-transcriptional regulation of gene expression is thought to rely on interactions between trans- and cis-acting factors^36^. In the context of tRNA modifications, the cis-acting factors have been assumed to consist of codons that require modified tRNAs for their decoding. For example, the *Dek* mRNA, which has a high frequency of U34-codons, was identified as a key gene showing high sensitivity to levels of U34-modified tRNA^15^. However, codons requiring U34-modified tRNA, although often referred to as rare, are present in the vast majority of mRNAs and scale with the length of the CDS (**Extended Fig. 5_01B)**. Therefore, if the hormone-dependent dynamic modulation of U34-modifications has a role in controlling cellular functions, it seemed plausible that additional features that do not simply scale with the length of the coding sequence would serve as cis-acting factors mediating sensitivity to U34-tRNA modifications. Consistently, our modeling of the ELP3-sensitive proteome indicated that frequencies of single U34 codons has a surprisingly small impact on transcript-specific regulation (**Fig. 3C**). Instead, we identified a limited set of di-codons, which we named ELP3 dependent down di-codons (“E3dDCs”) to reflect their role in controlling translation elongation, as a previously unrecognized mechanism whereby changes in U34 tRNA modifications remodel the cancer cell proteome to support proliferation in prostate cancer cells.

Importantly, our studies indicate that the presence of one of the six E3dDCs in the coding sequence is often insufficient to mediate sensitivity. This was explained by additional factors tuning the activity of E3dDCs including the local sequence context around E3dDCs as well as features in the 5’UTR affecting translation initiation. In aggregate, our findings reveal that E3dDCs are key components of coding sequence regions that act as translation traps with translation elongation dynamics sensitive to the availability of U34-modified tRNAs. Intriguingly, the sensitivity is also affected by the efficiency of initiation, whereby high initiation leads to a higher sensitivity to levels of U34-modified tRNA (**Fig. 6G**). A key interplay between elongation traps and initiation mechanisms is further supported by that mRNAs enriched in E3dDCs have reduced initiation efficiency (**Extended Fig. 5_01F**), a characteristic that seems to be conserved in human and mouse where E3dDCs-containing mRNAs harbor longer 5′UTRs (**Extended Fig. 5_01G**). Consequently, the coordination between initiation and elongation appears to establish a dynamic balance that determines the maximum protein synthesis rate from an mRNA. This mechanism may thereby also serve as a failsafe mechanism to limit the translation potential of a subset of mRNA as too high translation initiation would trigger elongation dysfunction and thereby restrict protein output. A similar failsafe mechanism acting at the level of mRNA translation has for example been observed for *AMD1* mRNA, where stop codon read through creates a translation “memory”, limiting the number of proteins that can be synthesized per mRNA molecule^37^.

The conserved enrichment of E3dDCs in cell-cycle–related genes supports their role in protecting cells from exaggerated protein synthesis with potential detrimental effects. As cancer cells often show augmented translation initiation of cell cycle related mRNAs^38^, this could explain the observed difference in sensitivity to reduced ELP3 expression between cancer and non-transformed cells’ proliferation. In principle, the translation trap is only triggered if translation initiation is too high relative to the elongation efficiency (**Fig. 6G**). This also indicates that there could be substantial variation in mRNAs that are affected depending on their efficiency of ribosome loading – thereby creating context dependence in sensitivity to reduced U34 tRNA-modifications. Indeed, the role of E3dDCs is not restricted to cell-cycle related genes as we show that despite induction of ISR, the associated expression program was not induced in a fashion paralleling frequencies of E3dDCs (**Fig. 4**). This role of ELP3 and U34 tRNA-modifications in regulating stress-sensitive gene expression is consistent with previous studies^39,40^ and the present study therefore provides a mechanism for how tRNA modifications controls stress-activated gene expression. As E3dDCs affect translation elongation whereas the acute ISR-program is induced via activation of translation initiation^30,31^, the results are consistent with downstream mechanisms affecting gene expression output in a dominant fashion. Indeed, variation in expression of mRNA does not necessarily correspond to alterations in protein expression due to downstream modulation of mRNA translation^41^. Our study therefore positions tRNA modifications as a central component during responses to cellular stress via enrichment of E3dDCs and points towards targeting of tRNA modifications in cancers addicted to stress-associated gene expression programs^42^.

A central challenge in interpreting cis-acting features regulating mRNA translation in the coding region, as opposed to for example RNA motifs in 5’UTR, is that they may reflect protein sequence rather than post-transcriptional regulation. However, multiple findings indicate that E3dDC function is independent of their encoded protein sequence including that: i) E3dDCs, but not the corresponding di-peptides, occur more frequent than what is expected from the frequencies of their individual codons/amino acids; ii) in contrast to U34-codons, E3dDCs frequencies do not parallel coding sequence length; and iii) E3dDCs are found in proteins with similar functions across species.

Here we report that reduced protein abundance upon ELP3 suppression is primarily caused by impaired protein synthesis rather than mRNA decay. A recent study^17^ linked m□A modifications within U34 codons as coordinating mRNA decay. Consistent with this, we also observed a negative correlation between reduced mRNA stability and higher frequency of U34-tRNA–decoded DRACH motifs within coding sequences. However, changes in mRNA stability only marginally accounted for the observed differences in protein levels (**Extended Figure 3_02C**). Our study therefore indicates that E3dDCs constitute the principal driver of the ELP3-sensitive proteome in prostate cancer. Yet, our data also support that these two programs are occurring in parallel. The lack of larger contribution of m□A in U34-tRNA–decoded DRACH is consistent with that there is no significant increase in the mcm□s²U/m□A ratio in prostate adenocarcinoma (PRAD) relative to their adjacent normal tissues^17^. Moreover, our quantification of m□A in the mRNA pool showed a trend for reduced levels of modifications **(extended Fig. 3_02E)**, which would limit the impact from this mechanism. Therefore, although both E3dDC- and m□A-dependent modulation of protein expression can occur in parallel downstream altered levels of U34-tRNA modifications, in prostate cancer, the E3dDC-dependent mechanism seems more pronounced.

So far, codon-pair effects have been investigated primarily in non-mammalian species. In yeast, it has been showed that translation speed is dictated not only by individual codons but also by adjacent codon pairs^28,43^. In mammalian systems, codon-pair de-optimization has been exploited to attenuate viral replication, providing functional evidence that codon-pair composition can influence protein output^44^. Complementary computational analyses have reinforced this concept, with CoCoPUTs and its cancer-focused extension CancerCoCoPUTs revealing non-random codon-pair usage and tumor type–specific signatures^45,46^. However, a role of codon-pairs in concert with their sequence context in generating translation traps sensitive to levels of tRNA modifications and efficiency of ribosome loading, as we identified herein, has to our knowledge not yet been reported.

A key component of our studies is application of the postNet algorithm^20^ that allows modeling and comparison of a multitude of mechanisms impacting post-transcriptional regulation in gene expression. In our study, it provided several key insights that would not have been possible using approaches that are presently regarded as state-of-the-art including: i) U34-codons and the reported hydrophilic short peptides, despite being enriched among mRNAs encoding proteins that were downregulated upon suppression of ELP3, only poorly explained observed changes in protein levels; ii) E3dDCs and E3uDCs explain ELP3-sensitive protein expression to an extent that U34-codons provide essentially no additional information; iii) it is the di-codons and not the associated di-amino acids that underlie observed regulation; iv) 5’UTR features affecting translation initiation explains ELP3-sensitive protein expression independently of features in the coding sequence; and v) the identified mechanisms depending on translation traps and initiation mechanisms is parallel and independent of the previous mechanism affecting mRNA stability depending on m□A in U34-tRNA–decoded DRACH motifs. This study thereby illustrates that efficient integration and modeling of cis-acting factors is essential to understand how post-transcriptional gene expression programs arise.

In summary, we present a model for how tRNA modifications modulate protein synthesis. In this model, di-codon context, rather than single-codon usage, underlies tunable elongation checkpoints whose impact on protein output is tuned by initiation kinetics. Furthermore, E3dDCs constitute key components that act as sensors for U34-modified tRNAs, whereby reduced levels of tRNA modifications can disrupt the balance between the ribosome loading and elongation capacity. Our data also support that precise tuning of initiation and elongation efficiency may have evolved as a mechanism restricting faulty hyperactive protein synthesis to e.g. oppose cancer development. Finally, prostate-cancer cells are highly dependent on U34-modified tRNA to sustain the balance between initiation and elongation efficiency, and thereby fuel their cancer-associated hyper-proliferation.

## MATERIALS AND METHODS

### Cell culture, organoids culture and proliferation assay

DU145, LNCaP and PNT1A cells were cultured in RPMI 1640 supplemented with 10% heat-inactivated FBS, 1% Penicillin/Streptomycin and 1X GlutaMAX in a humified incubator (5% CO_2_, 37°C). ELP3-depleted DU145, LNCaP and PNT1A cells were generated using CRISPR/Cas9 technology. To produce viral particles, HEK293T cells were transfected with plentiCRISPRv2 containing single guide RNA (sgRNA) targeting ELP3 (GATGCCTGACCTGCCAAACG (KO1) and GAGTTACTCTCCTAGTGACC (KO2); GenScript) and lentiviral packaging plasmids (pMDLg/pRRE, pRSV-Rev and pMD2.G) using Lipofectamine 3000 reagent (Invitrogen). DU145 cells were infected with viral particles and selected with puromycin. Single-cell clones were generated using two different sgRNAs and were expanded from single colonies obtained by sorting lentiviral-infected DU145 cells using a BD FACSAria™ Fusion5 (BD Biosciences), followed by immunoblotting screening. An empty vector (EV) control was generated for each cell line. For proliferation assays, cells from 6-cm plates were detached, stained with trypan blue and counted using a Countess II FL cell counter every 24 h.□

Organoids were derived from PDX tissues as previously described^22^. 201.1A and 305R organoids^22^ were seeded as droplets (1 x 10^5^ cells/droplet) in growth factor reduced, phenol red-free, ldEV-free Matrigel (Corning) and cultured in advanced DMEM/F-12 (Thermo Fisher) containing 1% penicillin-streptomycin, 2 mM Glutamax, 1 nM DHT, 1.25 mM N-acetylcysteine, 50 ng/ml EGF, 500 nM A83-01, 10 mM, nicotinamide, 10 μM SB202190 (Sigma), 2% B27 (Life Technologies), 100 ng/ml noggin (Peprotech), 10 ng/ml FGF10 (VWR), 5 ng/ml FGF2, 1 μM prostaglandin E2 (Tocris) and 10% R-spondin 1 conditioned media. 10 μM Y-27632 dihydrochloride (Selleck Chemicals) was added to culture media during organoid establishment and the following passage. Both organoid lines were infected with viral particles described above and selected with puromycin for 7 days followed by RT-qPCR analysis and PrestoBlue proliferation assay. For the PrestoBlue assay, media from each well was removed and replaced with 500 µL of a 1:10 dilution of PrestoBlue dye (Invitrogen) in DMEM/F12. Samples were incubated for 1 h in a humidified incubator (5% CO2, 37°C). After the incubation period, two 140 µL aliquots of the PrestoBlue dye were carefully removed from each well and transferred into a black, flat-bottomed 96-well plate (Corning), yielding 2 technical replicates per sample. Absorbance at 545 nm was measured using a Cytation 3 microplate reader (BioTek).

□

### Soft-agar colony formation assay

6-well plates were coated in 1.4% low melting point agarose solution prepared using culture media. The desired number of cells were prepared in equal volume of culture media and 0.7% low melting point agarose solution. 2 mL of the resulting 0.35% low melting point agarose-cell solution was layered on top of the 1.4% agar in each well. After solidification of the top layer, 100 µL of culture media was added to each well. 200 µL of complete growth media were added to each well every 3 days. After 3 weeks of incubation (37°C, 5% CO_2_), the cells were fixed and stained with 0.005% crystal violet solution for 1h at RT. The crystal violet solution was removed and the wells were washed with PBS. The plates were scanned using an Epson Perfection v850 Pro scanner and the images were converted to black and white prior to colony counting.

### tRNA high-performance liquid chromatography followed by tandem mass spectrometry

Small RNAs (<200 nucleotides) were isolated using the mirVana Kit (Invitrogen AM1516) following the manufacturer’s protocol for the “enrichment procedure for small RNA” protocol. 4 μl of small RNA (383-776.5 ng) was used to evaluate levels of cm^5^U, mcm^5^U, and mcm^5^s^2^U by LC-MS/MS as described in Tardu et al, 2019^47^. Each sample was mixed with 0.4 pg/μl of isotopically labeled guanosine ([^13^C][^15^N]-G) as the internal standard. The enzymatic digestion was carried out using Nucleoside Digestion Mix (New England BioLabs) according to the manufacturer’s protocol. The digested samples were lyophilized and reconstituted in 100 μl of RNAse-free water containing 0.01% formic acid prior to UHPLC-MS/MS analysis. The UHPLC-MS analysis was performed on a Waters XEVO TQ-S™ (Waters Corporation, USA) triple quadrupole mass spectrometer equipped with an electrospray ionization source (ESI) maintained at 150°C and a capillary voltage of 1 kV. Nitrogen was used as the nebulizer gas which was maintained at 7 bars pressure with a flow rate of 500 l/h at 500°C. UHPLC-MS/MS analysis was performed in ESI positive-ion mode using multiple reaction monitoring (MRM) from ion transitions previously determined for mcm^5^s^2^U. A Waters ACQUITY UPLC™ HSS T3 guard column, 2.1 × 5 mm, 1.8 μm, attached to an HSS T3 column, 2.1 × 50 mm, 1.7 μm was used for the separation. Mobile phases included RNAse-free water (18 MΩ cm−1) containing 0.01% formic acid (Buffer A) and 50:50 acetonitrile in Buffer A (Buffer B). The digested nucleotides were eluted at a flow rate of 0.5 ml/min with a gradient as follows: 0–2 min, 0–10% B; 2–3 min, 10–15% B; 3–4 min, 15–100% B; 4–4.5 min, 100% B. The total run time was 7 min. The column oven temperature was kept at 35°C, and sample injection volume was 10 μl. Three injections were performed for each sample. Data acquisition and analysis were performed with MassLynx V4.1 and TargetLynx. Quantification was performed based on nucleoside-to-base ion transitions using calibration curves generated from pure nucleoside standards and the stable isotope-labeled guanosine IS. mcm^5^U and mcm^5^s^2^U were synthesized by Dr. Malkiewicz and colleagues, and cm^5^U was purchased from the AA BLOCKS LLC. (San Diego, CA, USA). [^13^C][^15^N]-G was purchased from the Cambridge Isotope Laboratories, Inc. (Tewksbury, MA, USA). All the nucleoside standards were characterized individually by liquid chromatography coupled with both UV and mass spectrometric detection.

### Western blotting

Protein lysates were prepared using radioimmunoprecipitation assay (RIPA) buffer supplemented with protease inhibitor cocktail (Roche), phenylmethylsulfonyl fluoride (PMSF; Thermo Scientific) and sodium orthovanadate (Na_3_VO_4_; NEB) (Supp. Table 1). Protein concentrations were determined using the Pierce bicinchoninic acid (BCA) assay (Thermo Scientific). Protein lysates were mixed with 6X protein sample loading buffer (Supp. Table 1) and resolved by SDS-PAGE on hand-cast polyacrylamide gels containing 8% or 10% acrylamide/bisacrylamide. Gels were run using Mini-Protean Tetra Cell system (Bio-Rad) in Tris-Glycine-SDS running buffer (Supp. Table 1) at 100 V for 2 h at room temperature (RT). Proteins were transferred from polyacrylamide gels to polyvinylidene difluoride (PVDF) membranes (Millipore) in Tris-glycine-methanol transfer buffer (Supp. Table 1) at 250 mA for 1.5 h at 4°C. Membranes were blocked for 1 h at RT and incubated overnight at 4°C with primary antibody, prepared with antibody-specific dilutions (Supp. Table 2) in primary antibody diluent solution (Supp. Table 1). The following day, membranes were washed 3 times in Tris buffered saline supplemented with 0.1% Tween-20 (TBS-T) (Supp. Table 1) and incubated with the appropriate HRP–conjugated secondary antibody (Supp. Table 2), diluted 1:5000 in blocking buffer (Supp. Table 1) for 1 h at RT. Membranes were washed 3 times in TBS-T and incubated for 1 min in Clarity Western enhanced chemiluminescence (ECL) Substrate (Bio-Rad) and protein signals were visualized using a ChemiDoc XRS+ imaging system (Bio-Rad). Signals were quantified using ImageJ software, measuring band intensities relative to β-actin control.□

**Supplementary Table 1.** List of reagents used in immunoblotting analysis.

| Reagent/Buffer | Recipe |
| --- | --- |
| Radioimmunoprecipitation assay buffer (RIPA) | 10 mM Tris-HCl, pH 8.0, 1 mM EDTA, 0.5 mM EGTA, 1% Triton X-100, 0.1% Sodium Deoxycholate, 0.1% SDS, 140 mM NaCl |
| 6x Protein sample loading buffer | 2% (w/v) SDS, 0.4 M Tris-HCl pH 6.8, 48% (v/v) Glycerol, 58 mM β-mercaptoethanol, 0.25% (w/v) Bromophenol Blue |
| Tris-Glycine-SDS running buffer | 25 mM Tris, 192 mM Glycine, 0.1% (w/v) SDS pH 8.7 |
| Tris-Glycine-methanol transfer buffer | 25 mM Tris, 192 mM Glycine, 15% (w/v) methanol |
| Tris buffered saline (TBS) | 50 mM Tris, 150 mM NaCl |
| Blocking buffer | 5% (w/v) skim milk in TBST (0.1% (v/v) Tween-20 in TBS) |
| Primary antibody diluent solution | 3% bovine serum albumin, 0.01% sodium azide |

**Supplementary Table 2.**
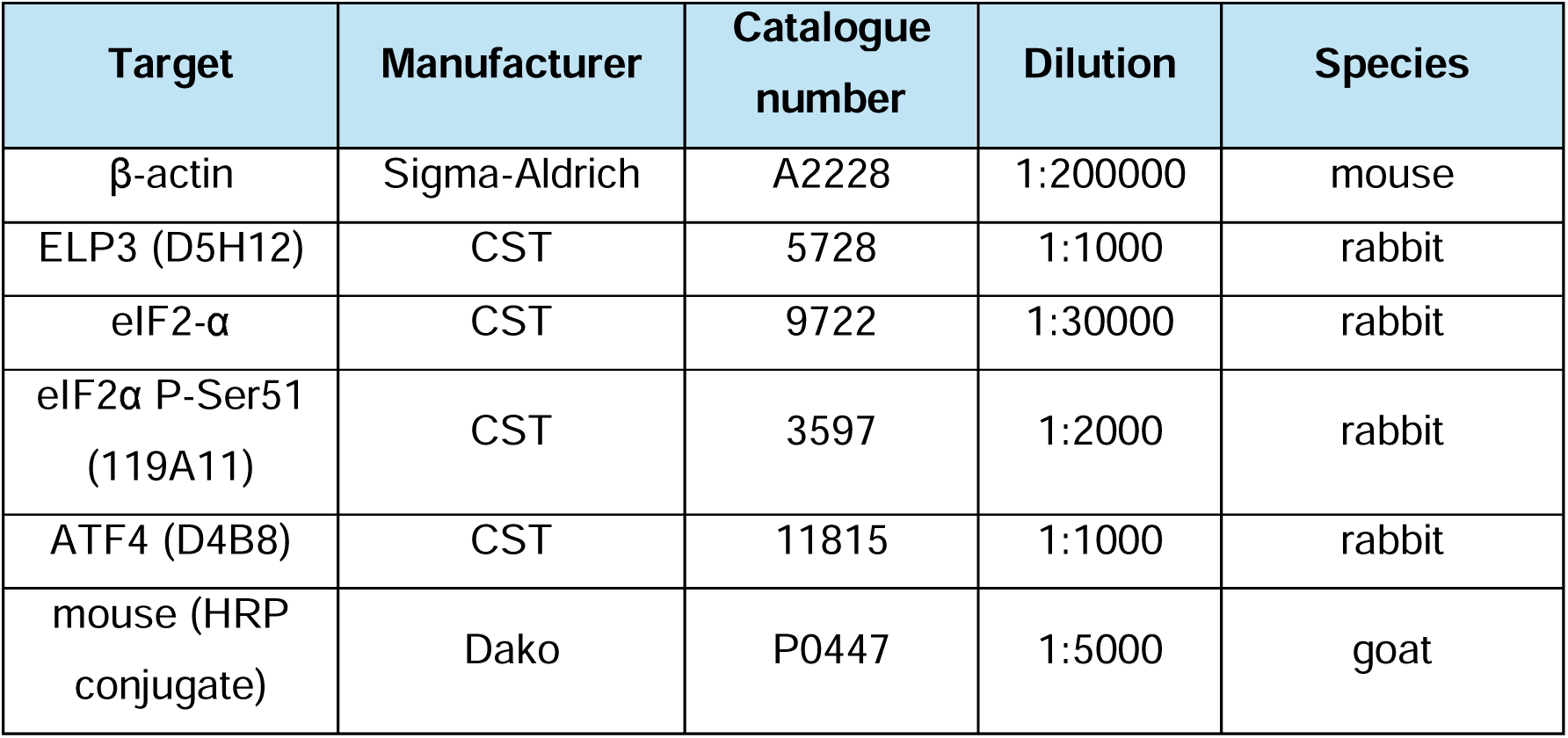
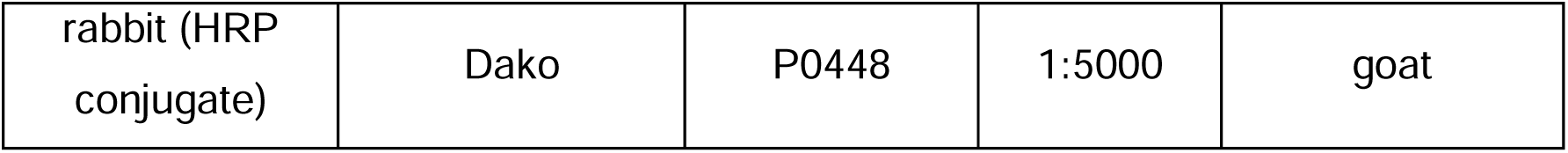
List of antibodies used in immunoblotting analysis.

| Target | Manufacturer | Catalogue number | Dilution | Species |
| --- | --- | --- | --- | --- |
| β-actin | Sigma-Aldrich | A2228 | 1:200000 | mouse |
| ELP3 (D5H12) | CST | 5728 | 1:1000 | rabbit |
| eIF2-α | CST | 9722 | 1:30000 | rabbit |
| eIF2α P-Ser51 (119A11) | CST | 3597 | 1:2000 | rabbit |
| ATF4 (D4B8) | CST | 11815 | 1:1000 | rabbit |
| mouse (HRP conjugate) | Dako | P0447 | 1:5000 | goat |
| rabbit (HRP conjugate) | Dako | P0448 | 1:5000 | goat |

### Live cell imaging

10,000 cells were plated in μ-slide 8-well chamber slides (Ibidi 80826) onto 250 μL of media at 24 hr prior to imaging on Olympus FV 3000 confocal microscope. Cells were stained with SPY-650-DNA stain (1 in 1000, Spirochrome, SC501) for 1 hr before imaging according to manufacturer’s instruction. Live cell time-lapse microscopy was performed on Plapon 60x OSC2 oil immersion objective for 18 hr with pictures taken every 8 mins. Laser 635nm was used on Alexa flour channel. 2 positions per well were captured. Data were analyzed using Fiji Software.

### Reverse transcription quantitative real-time polymerase chain reaction (RT-qPCR)

Total RNA was extracted using the RNeasy Mini Kit (Qiagen) according to the manufacturer’s protocol. RNA yield and purity were assessed with a Nanodrop ND1000 spectrophotometer (Thermo Fisher Scientific). Equal amounts of RNA were treated with RNase-free DNase (Promega) at 37°C for 15 min, followed by enzyme inactivation at 70°C for 15 min. Complementary DNA (cDNA) was generated using 60 U Superscript III reverse transcriptase (Life Technologies) and 12.25 ng random hexamer primers (Promega), with the following thermal profile: 5 min at room temperature, 5 min at 37°C, 120 min at 47°C, and 15 min at 70°C. The resulting cDNA was diluted in nuclease-free water and stored at −20°C. For quantitative real-time PCR, reactions consisted of 8 μl cDNA sample and 12 μl Fast SYBR Green Master Mix (Applied Biosystems) containing 0.1 μM forward and reverse primers (Supp. Table 3), in triplicate wells of MicroAMP Optical 96-well plates (Applied Biosystems). Amplification was performed on a StepOne Plus Real-Time PCR system (Applied Biosystems,). Relative expression levels were calculated using the 2^−ΔΔCt^ method.

**Supplementary Table 3.**
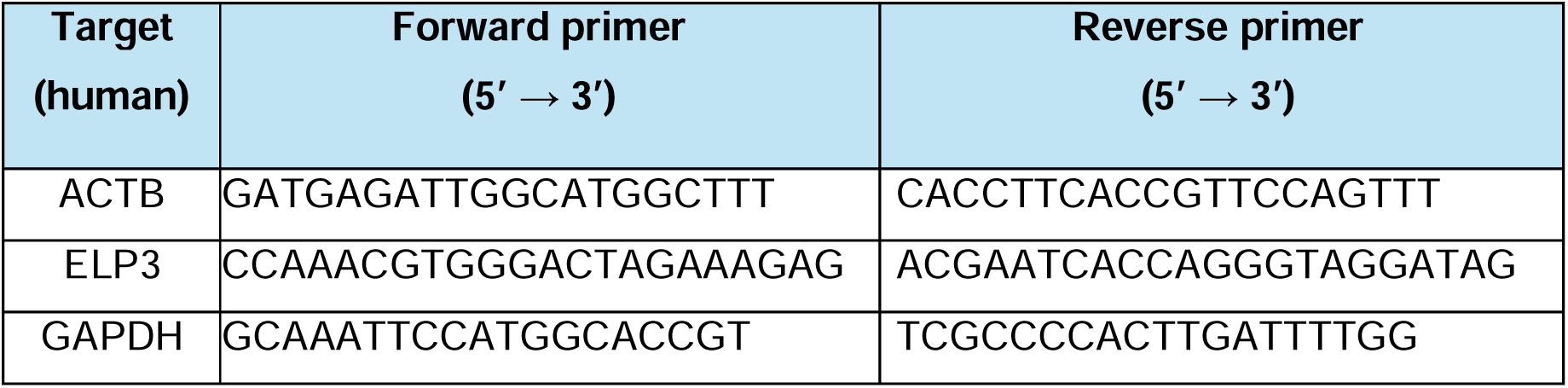
List of RT-qPCR primers.

### m□A quantification in poly(A)+ RNA

Total RNA was extracted from DU145^EV^, DU145^KO1^, and DU145^KO2^ cells using the RNeasy Mini Kit (Qiagen 74106) according to the manufacturer’s instructions. Total RNA concentration and purity were evaluated using a NanoDrop ND-1000 spectrophotometer (Thermo Fisher Scientific). Poly(A)+ RNA was isolated from 2.25 µg of total RNA per sample using the NEBNext®− Poly(A) mRNA Magnetic Isolation Module (New England Biolabs E7490S). The enrichment and concentration of poly(A)+ RNA samples were assessed with the Agilent TapeStation 4150 using High Sensitivity RNA ScreenTape (Agilent Technologies). Equal amounts of poly(A)+ RNA from each cell line were subjected to quantification of m□A levels using the m□A RNA Methylation Assay Kit (Abcam ab185912) following the manufacturer’s protocol. Absorbance at 450 nm was measured using a Cytation3 plate reader (BioTek).

### Label-free quantification (LFQ) proteomics

Liquid chromatography coupled with tandem mass spectrometry (LC-MS/MS) was performed on four biological replicates for each cell line. For each replicate, cells from one T75 flask were washed with PBS, trypsinised for 3 min in a humidified incubator (37°C, 5% CO₂), and collected in a 15 mL centrifuge tube. The cells were centrifuged at 250 RCF for 3 min at room temperature (RT), and the resulting pellet was resuspended in 5 mL PBS -/-. Samples were centrifuged again at 250 RCF for 3 min at RT, and the pellets were transferred to 1.5 mL microcentrifuge tubes and resuspended in 1 mL PBS -/-. After a final centrifugation at 250 RCF for 3 min at RT, the supernatant was carefully removed. The pellets were snap-frozen in liquid nitrogen (LN₂) and stored at –80°C until further processing. Cell pellets were solubilised in 5% SDS–100 mM Tris-HCl by heating at 95°C for 10 min to denature endogenous proteases and phosphatases. DNA was sheared using probe sonication. Samples were processed using the S-Trap protocol according to the manufacturer’s instructions^48^. Eluted peptides were acidified to 1% trifluoroacetic acid and purified using StageTips packed with SDB-RPS (Empore)^49^. Indexed retention time (iRT) standard peptides were spiked into all samples prior to LC-MS/MS analysis for quality control^50^. Samples were processed using a Dionex Ultimate 3000 RSLC Nano and Orbitrap Exploris 480 mass spectrometer (Thermo Scientific) using Acclaim PepMap RSLC (75 μm x 50 cm, nanoViper, C18, 2 μm, 100Å) analytical column (Thermo Scientific).□

Raw data files from DU145^EV^, DU145^KO1^ and DU145^KO2^ cells were processed using MaxQuant v2.0.2.0 employing the built-in MaxLFQ algorithm^51,52^ to identify peptides and their label-free quantification (LFQ) values using in-house standard parameters. MaxLFQ calculates normalized protein intensities from peptide-level signals across all runs. For downstream analyses, only the LFQ intensity values were used as a proxy for relative protein abundance. Proteins with Q-value < 0.05 were retained, and those with missing or zero LFQ values in any sample were excluded, yielding a complete dataset across all conditions. LFQ intensities were log₂-transformed prior to statistical analysis. Data reproducibility across EV control and knockout samples was assessed by principal component analysis (PCA) after filtering proteins by variance, retaining only those in the top quartile of standard deviation across samples. Changes in protein abundance relative to the empty vector control were analyzed using the anota2seq algorithm^24^. As anota2seq requires a matched input, a mock dataset was generated to enable protein-only analysis. Quality control indicated no evidence of batch effects; thus, no batch correction was applied. Significance thresholds were set at maxRvmPAdj = 0.05 (corresponding to FDR < 5%).

### Gene-ontology enrichment analysis

Gene ontology (GO) enrichment analysis was performed using the enrichGO function from the clusterProfiler^53^ R package on the set of proteins identified by anota2seq as significantly regulated. The background was defined as all proteins passing the thresholds described above for anota2seq input. Analyses were conducted for the Biological Process (BP) ontology using org.Hs.eg.db as the annotation database, with keyType = “SYMBOL”. Parameters were set to pAdjustMethod = “BH”, pvalueCutoff = 0.05, qvalueCutoff = 0.05, minGSSize = 10, and maxGSSize = 300. Enriched GO terms were simplified to remove redundancy using the simplify function with the Wang semantic similarity method (measure = “Wang”, cutoff = 0.6).

### Cross-species GO-analysis

For human and mouse, the longest CDS isoform per gene from the CCDS database was used to quantify E3dDC content, and genes were grouped according to the number of E3dDC present in their CDS (no-E3dDC and multi-E3dDC). Within each species, GO-BP enrichment was performed separately for each group using org.Hs.eg.db (human) and org.Mm.eg.db (mouse), with the same clusterProfiler parameters as described above. The background universe for each species was defined as all genes included in that species’ dataset. For yeast, canonical coding sequences of Saccharomyces cerevisiae S288C were obtained from the Saccharomyces Genome Database (SGD; orf_coding.fasta.gz, http://sgd-archive.yeastgenome.org/sequence/S288C_reference/orf_dna/orf_coding.fasta.gz), and genes were grouped by E3dDC count using the same approach. Enrichment analysis was carried out with org.Sc.sgd.db (keyType = “GENENAME”) using the same multiple-testing settings, with the background defined as all verified/standard ORFs from the SGD CDS set. Cross-species comparisons were then carried out separately for the multi-E3dDC and no-E3dDC groups. In the multi-E3dDC group, enriched GO terms were merged across species by GO identifier, and terms were retained only if they appeared in all three species and were significant at FDR < 0.05 in each. In the no-E3dDC group, enriched terms were likewise merged across species, and terms were retained if they were present in all three species and significant in at least one (FDR < 0.01). For visualization, GO terms were ordered by their significance in the human dataset, with the corresponding enrichment statistics from mouse and yeast were arranged in the same order.

### Polysome Profiling

Cells were treated with 100 µg/mL cycloheximide for 5 min at 37°C prior to harvesting. Following incubation, the culture media was removed and cells were washed with 1X hypotonic wash buffer (Supp. Table 4). Cells were harvested in 1X hypotonic lysis buffer (Supp. Table 4) using a sterile scraper, and the lysates were transferred to new sterile, nuclease-free 1.5 mL tubes. Samples were centrifuged at 16,000 RCF for 7 min at 4°C, and the supernatant was collected into sterile, nuclease-free 2 mL tubes. 10% of each lysate was transferred into a fresh sterile, nuclease-free 2 mL tube containing 860 µL TRIzol reagent (Invitrogen), mixed thoroughly, and stored at –80°C for later use as “input cytoplasmic mRNA” controls. The remaining lysates were loaded onto a 10–40% sucrose density gradient, ultracentrifuged for 2 h 15 min at 36,000 RPM (222,228 RCF) at 4°C, and fractionated with a Foxy Jr. Fraction Collector at a rate of one fraction per minute, yielding a final volume of 860 µL per fraction. During fractionation, absorbance at 260 nm was monitored in real time using an ISCO UA-6 UV/Vis detector. Fractions were collected, mixed with 860 µL TRIzol (Invitrogen), and stored at –80°C until further use. Cytoplasmic and polysome-associated RNA were extracted from the TRIzol-mixed lysates according to the manufacturer’s instructions and further purified using the Qiagen RNeasy Mini Kit (Qiagen 74106). RNA concentrations were determined with a NanoDrop ND-1000 spectrophotometer (Thermo Fisher Scientific), and integrity was assessed using the Agilent 2100 TapeStation System. Only samples with RIN >8 were used for library preparation. Samples were stored at –80°C until further use. For library preparation, rRNA was depleted using the NEBNext® rRNA Depletion Kit (Human/Mouse/Rat, v1; New England Biolabs E6310) according to the manufacturer’s protocol. The resulting rRNA-depleted RNA was used as input for library construction with the NEBNext® Ultra™ II DNA Library Prep Kit for Illumina (NEB #E7103). Libraries were sequenced on the Illumina NextSeq 500 platform (single-end, 75 bp reads), generating ∼40 million reads per sample.

**Supplementary Table 4.** Components of buffers used in polysome profiling analysis.

| Solution | Recipe |  | Supplier |
| --- | --- | --- | --- |
|  | Component | Final concentration |  |
| 10X<br>hypotonic□<br>buffer□ | Tris-HCl pH 7.5 | 0.05 M | Sigma-Aldrich T3253 |
|  | MgCl<sub>2</sub> | 0.025 M | Sigma-Aldrich M4880 |
|  | KCl | 0.015 M | Sigma-Aldrich P9333 |
| 1X Hypotonic wash buffer | 10X hypotonic buffer | 1X | See above |
|  | cychloheximide | 100 µg/mL | Sigma-Aldrich C1988 |
| 1X Hypotonic lysis buffer | 10X hypotonic buffer | 1X | See above |
|  | Sodium deoxycholate | 0.50% | Sigma-Aldrich D6750 |
|  | Triton X-100 | 0.50% | Sigma-Aldrich T8787 |
|  | cOmplete, EDTA-free protease inhibitor cocktail | 1X | Roche 4693132001 |
|  | RNasin | 10 U | Promega N2111 |
|  | cychloheximide | 100 µg/mL | Sigma-Aldrich C1988 |
|  | 1,4-dithiothreitol (DTT) | 2 mM | Sigma-Aldrich 43816 |

### RNA-seq data preprocessing and analysis

RNA sequencing data were processed using the nf-core/rnaseq pipeline^54^ under Singularity. Reads were aligned to the human reference genome GRCh38 using HISAT2^55^, with default quality trimming and alignment settings. Coordinate-sorted BAM files produced by the pipeline were exported, and gene-level quantification was performed independently in R using featureCounts^56^ with the following settings: isGTFAnnotationFile = TRUE, single-end (isPairedEnd = FALSE), reverse stranded (strandSpecific = 2), exon features (GTF.featureType = “exon”), excluding multi-mapping reads (countMultiMappingReads = FALSE) and junction counts (juncCounts = FALSE), fraction = FALSE. Prior to counting, the RefSeq GTF file (GCF_000001405.40, GRCh38.p14) was pre-processed by converting chromosome names to the “chr” format, removing scaffold and alternative contigs, and retaining only standard chromosomes (chr1–22, chrX, chrY).

Prior to statistical analysis, genes with zero read counts in any sample were removed. Genes were then further excluded if, in the empty vector control group, any replicate had fewer than 100 reads. The remaining protein-coding genes were normalized using the trimmed mean of M values (TMM) method^57^. Normalized counts were then transformed to log₂ scale with the voom function within the anota2seq R package^24^. To detect changes in translation, normalized counts from polysome-associated and cytosolic mRNA fractions were analyzed using anota2seq algorithm with the following slope and delta cutoffs (deltaPT = deltaTP = log₂(1.2), deltaP = deltaT = 0, minSlopeTranslation = -1, maxSlopeTranslation = 2, minSlopeBuffering = -2, maxSlopeBuffering = 1). The resulting p-values were adjusted for multiple testing using the Benjamini-Hochberg procedure, and FDR < 15% (by setting maxRvmPAdj < 0.15) was considered significant. Gene expression alterations were classified into three categories, altered mRNA abundance, changes in translation efficiency, or translational offsetting via the anota2seqRegModes function.

### Modeling regulated proteins with “postNet”

Protein fold-changes for regulated proteins identified by anota2seq were used as input for modeling. Modeling was performed using the featureIntegration function from postNet with the following parameters: analysis_type = “lm”, regulationGen = “translatedmRNA” (= regulated proteins), regOnly = TRUE, allFeat = FALSE, useCorel = TRUE, and NetModelSel = “omnibus”. The modeling workflow begins with univariate linear models, in which each variable is tested for its association with the measured effect (effM). The variable explaining the largest proportion of regulation is then used to initiate a stepwise regression. Additional variables are sequentially added to the model. In the final step, correlations among the selected variables are adjusted for all other included variables to estimate the adjusted contribution of each feature to the explained regulation. Unless otherwise specified, statistical significance was assessed at an FDR threshold of 0.05, as implemented by the package.

To define sets of cis-acting variables, we used the longest CDS isoforms from the CCDS database and their corresponding UTR regions from RefSeq. For Model 1, CDS-derived variables included GC3% content, codons decoded by mcm^5^s^2^, mcm^5^, and cm^5^ U34-modified tRNA, and CDS length. Experimentally supported RNA-binding protein (RBP) motifs were obtained from the ATtRACT database^58^, and quantified in the CDS of the transcripts corresponding to the regulated proteins. Additional CDS-level variables included pentamer motifs^19^, U34-DRACH, GAA-DRACH, and AGA-DRACH motifs.

De novo motifs were identified with MEME algorithm from MEME Suite^59^ in differential enrichment mode using CDS of downregulated proteins as the primary set and non-regulated CDS as control. A first-order Markov background model was generated from the non-regulated sequences using fasta-get-markov (parameters: - m 1 -rna -norc). The codon usage indexes were computed with codonW (https://sourceforge.net/projects/codonw/files/codonw/SourceCode-1.4.4%28gz%29/) and tAI packages^60^. For UTR-features, folding energy was calculated using the Mfold algorithm^61^. G-quadruplexes were predicted with pqsfinder (min_score = 47)^62^. The poly(A) tail lengths were obtained from published datasets^63,64^.

### Exon-Intron Split Analysis (EISA)

EISA was performed to distinguish transcriptional from post-transcriptional changes in gene expression^27^. Single-end RNA-seq reads were first aligned to the human reference genome GRCh38 using HISAT2^55^. Exon and intron annotation ranges were obtained from the RefSeq GTF (GCF_000001405.40, GRCh38.p14) using the getFeatureRanges() function in eisaR pacakge with parameters featureType = c(“spliced”,“intron”), intronType = “separate”, flankLength = 90L, joinOverlappingIntrons = FALSE, collapseIntronsByGene = FALSE.

Read counting was performed using featureCounts^56^ (Rsubread package), with exon and intron features quantified separately against custom SAF annotation files. Parameters were set as follows: isGTFAnnotationFile = FALSE, isPairedEnd = FALSE, strandSpecific = 2, countMultiMappingReads = FALSE, fraction = FALSE, juncCounts = FALSE, useMetaFeatures = TRUE, allowMultiOverlap = FALSE. Counts from the two sequencing lanes were summed per gene. To ensure comparability, only genes with at least one nonzero count in both exonic and intronic fractions were retained. Intronic gene identifiers were standardized to match exon identifiers, as required by the EISA framework to ensure proper pairing of exon–intron counts. Filtered exon and intron count matrices were then provided as input to the runEISA() function (Version 1.21.0), with the following parameters: modelSamples = TRUE, geneSelection = “filterByExpr”, statFramework = “QLF”, effects = “predFC”, recalcNormFactAfterFilt = TRUE. All other arguments were left at default settings.

### Signature sets used for ISR analysis

Signatures of translation regulation by ISR pathway were obtained from published studies or from re-analysis of published datasets using the anota2seq algorithm to define sets of genes translationally activated or inhibited by ISR^30,31^. These gene sets were then used as signatures for downstream analyses in our study. To evaluate the relative behavior of ISR signature genes, we applied empirical cumulative distribution function (ECDF) analysis, which plots the cumulative proportion of genes as a function of their fold-change values. This allowed direct comparison of the distribution of fold-changes for ISR signatures against background genes.

### Local feature analysis

To investigate the local sequence context of E3dDCs, we extracted 102-nucleotide regions surrounding of each E3dDC within the coding sequence of the transcripts. The E3dDC was excluded from the windows. The resulting sequences were used to obtain cis-acting feature variables for downstream analyses. Features were aggregated at the gene level by assigning the extreme value (maximum or minimum, depending on the biological interpretation of the feature) across all 102-nucleotide regions as the representative value for each gene in downstream modelling. Statistically significant features within each of the following groups: individual amino acids, codons, di-codons and di-peptides, were retained (FDR < 0.15) for inclusion in the final model explaining the regulation of multi-E3dDC proteins. For 5’UTR features, human RNA-binding protein (RBP) motifs were obtained from the ATtRACT database^58^. Motifs shorter than six nucleotides were excluded and motifs were required to be present in at least 1% of the regulated proteins. Features with FDR < 0.15 were retained for inclusion in the final model. To capture nascent peptide context, 102-nucleode regions were analyzed for hydrophobicity. Hydrophobicity was computed using the Kyte–Doolittle scale^65^ (Peptides R package 2.4.6). All candidate features were pooled, and the final model was based on an FDR threshold of 0.15.

### Dual-fluorescence assay by flow cytometry

Plasmids for dual-fluorescence assays were generated by inserting linker sequences, including poly-lysine (21×A repeat), STAG2, and TRMT6 gene fragments, between GFP and mCherry (ChFP) coding sequences, as described previously^34^. Fragments from STAG2 (CCDS43990; fragment length= 432nt) and TRMT6 (CCDS13093, fragment length= 231nt) were obtained from the CCDS database and correspond to specific regions within their CDSs (STAG2: E3dDC region starting at nt 2494 and control region at nt 2059; TRMT6: E3dDC region at nt 277 and control region at nt 754). DU145^EV^ and DU145^KO1^ cells were transfected with the appropriate dual-fluorescence constructs using Lipofectamine 3000 reagent (Thermo Fisher Scientific). Cells were incubated for 18 h in transfection media containing Lipofectamine, followed by replacement with complete growth media and an additional 18 h incubation prior to harvesting. Cells were analyzed on a BD FACSymphony™ A3 Cell Analyzer (BD Biosciences) using BD FACSDiva™ software. Spectral overlap between fluorophores was compensated using single-stained controls.

For each biological replicate, FCS files were imported into R using the read.flowSet() function and subsequently analyzed, with unstained control samples used to define the gating coordinates. Cells were first gated on FSC-A versus SSC-A with an elliptical gate to exclude debris. Singlets were then selected using a diagonal polygon gate on FSC-A versus FSC-H, and live cells were retained by excluding events with high signal in the near-infrared viability dye channel (threshold = 25; Invitrogen L34976). The gates derived from unstained controls were then consistently applied across GFP-, mCherry-, and experimental samples. Fluorescence signals for GFP and mCherry were extracted and transformed using a logicle transformation^66^ (flowCore::estimateLogicle) to stabilize variance. GFP-positive cells were defined as those with transformed GFP intensity greater than one. To control for technical variation, cells expressing the linker (efficient translation) and polyK (inefficient translation) constructs were isolated. mCherry signal was then adjusted using a linear regression model with GFP intensity, replicate, and cell type as covariates, and residuals were extracted. Cells were subsequently stratified into quartiles based on GFP intensity, and the density distributions of residuals were compared between linker and polyK within each quartile. This analysis revealed that the separation between linker and polyK progressively increased across quartiles, with the strongest differences observed in the upper two quartiles (Extended Fig. 6_01A). For experimental constructs (i.e., STAG2 and TRMT6), mCherry signal was adjusted using the same linear regression model (mCherry ∼ GFP + replicate + cellType), and residuals were extracted. Residuals from the two highest GFP-expression quartiles were pooled and visualized using ggplot2. To assess whether translation effects associated with the constructs varied depending on the cell type, we fitted a linear model with the design:

*mCherry_ijk_ ∼ b_0_ + b_1_.GFP_ijk_ + b_2_.replicate_i_ + b_3_.construct_j_ + b_4_.cellType_k_ + b_5_.(constrcut_j_ * cellType_k_) + ε_ijk_*

where Y is the mCherry fluorescence intensity measured in replicate i, construct j, and cellType k; *GFP_ijk_*is the corresponding GFP intensity; *replicate_i_* accounts for biological replicate effects; *construct_j_* denotes the expressed region of interest; *cellType_k_* indicates the cell line; and *b_5_.(constrcut_j_ * cellType_k_)* is the interaction term testing whether the effect of region differ depending on the cell type. ε*_ijk_* represents the residual error.

### Statistical analysis

Statistical tests, significance thresholds, and quantification approaches are described in detail in the relevant Methods subsections or in the corresponding figure legends.

## DATA AVAILABILITY

Processed RNA-seq data and metadata are available in BioStudies under accession S-BSST2300 and raw RNA-seq data have been deposited in the European Nucleotide Archive (ENA) under accession PRJEB104146. Proteomics data have been deposited in PRIDE under accession PXD071176. In the deposited raw files, the sample identifiers are as follows: DU145^KO1^ = 1A2, DU145^KO2^ = 4A3.

## ACKNOWLEDGEMENTS

The authors thank Q. Lin for nucleoside mass spectrometry. The authors also acknowledge the Molecular Genomics Core (RRID:SCR_025695), the Research Flow Core (RRID:SCR_025550) and the Centre for Advanced Histology and Microscopy (RRID:SCR_025432) at the Peter MacCallum Cancer Centre for access and support. This research was supported by a National Health and Medical Research Council (NHMRC) Ideas Grant (2019032) to LF, EPK and OL; grants from Swedish Research Council (2020-01665), Swedish Cancer Society (222186), the cancer research funds of Radiumhemmet (231263), and a Wallenberg Academy Fellowship to OL; a Victorian Cancer Agency Early Career Fellowship (ECRF20018) and Prostate Cancer Foundation of Australia Future Leaders Fellowship 2024 to EPK.

## AUTHOR CONTRIBUTIONS

EPK, LF and OL conceived the study and designed the experiments. CT, EPK, LNG, CAT and JRS performed experiments. KHM and OL performed bioinformatic data analysis. KHM, CT, JRS, RBS, EPK, LF and OL analyzed and interpreted the data. EPK, LF and OL supervised the study. KHM, EPK, LF and OL wrote the manuscript with input from all authors.

## EXTENDED FIGURE LEGENDS

**extendedFigure2_01:**
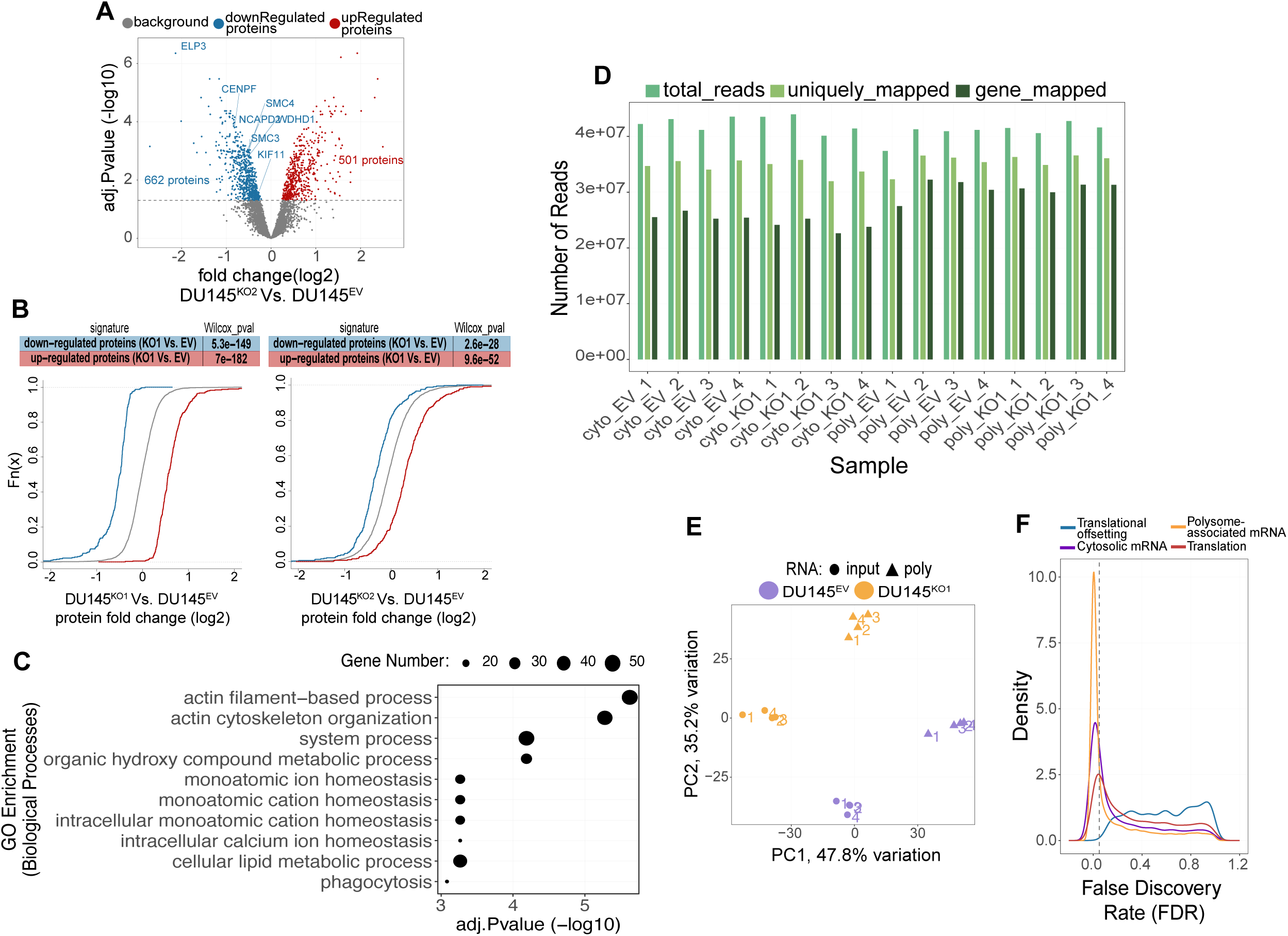
ELP3 suppression blocks elongation of cell-cycle related proteins. A) Volcano plot for changes the in proteome between DU145^KO2^ and DU145^EV^ cells. ELP3 and the number of regulated proteins are indicated. B) Empirical cumulative distribution functions (log2 fold changes) for the set of proteins identified as regulated in DU145^KO1^ vs DU145^EV^ cells in DU145^KO1^ Vs. DU145^EV^ (left) and DU145^KO2^ Vs. DU145^EV^ (right). Grey lines show the corresponding background distributions (all quantified proteins). Differences between sets of regulated proteins and the background were assessed using a two-sided Wilcoxon rank-sum test and the obtained p-values are indicated in the table above the plots. C) GO enrichment analysis of upregulated proteins following ELP3 suppression (DU145^KO1^ Vs. DU145^EV^). Displayed are significantly enriched biological processes with an FDR < 0.001. D) Number of RNA-seq reads from polysome-profiling across samples and processing steps. Bars represent total sequenced reads, uniquely mapped reads, and reads assigned to genes (left to right) for cytoplasmic (cyto) and polysome-associated (poly) RNA fractions from DU145 empty vector (DU145^EV^) and ELP3 knockout (DU145^KO1^) cells (n = 4). E) PCA of per-gene centered expression values from polysome profiling (N = 4), showing the first two principal components. F) Kernel densities of false discovery rates (FDR) for differential expression in DU145^KO1^ versus DU145^EV^. Shown are results from analysis of polysome-associated mRNA (orange), cytosolic mRNA (purple), translation (i.e. polysome-associated mRNA changes after adjustment for total mRNA changes; red), and translational offsetting (blue).

**extendedFigure3_01:**
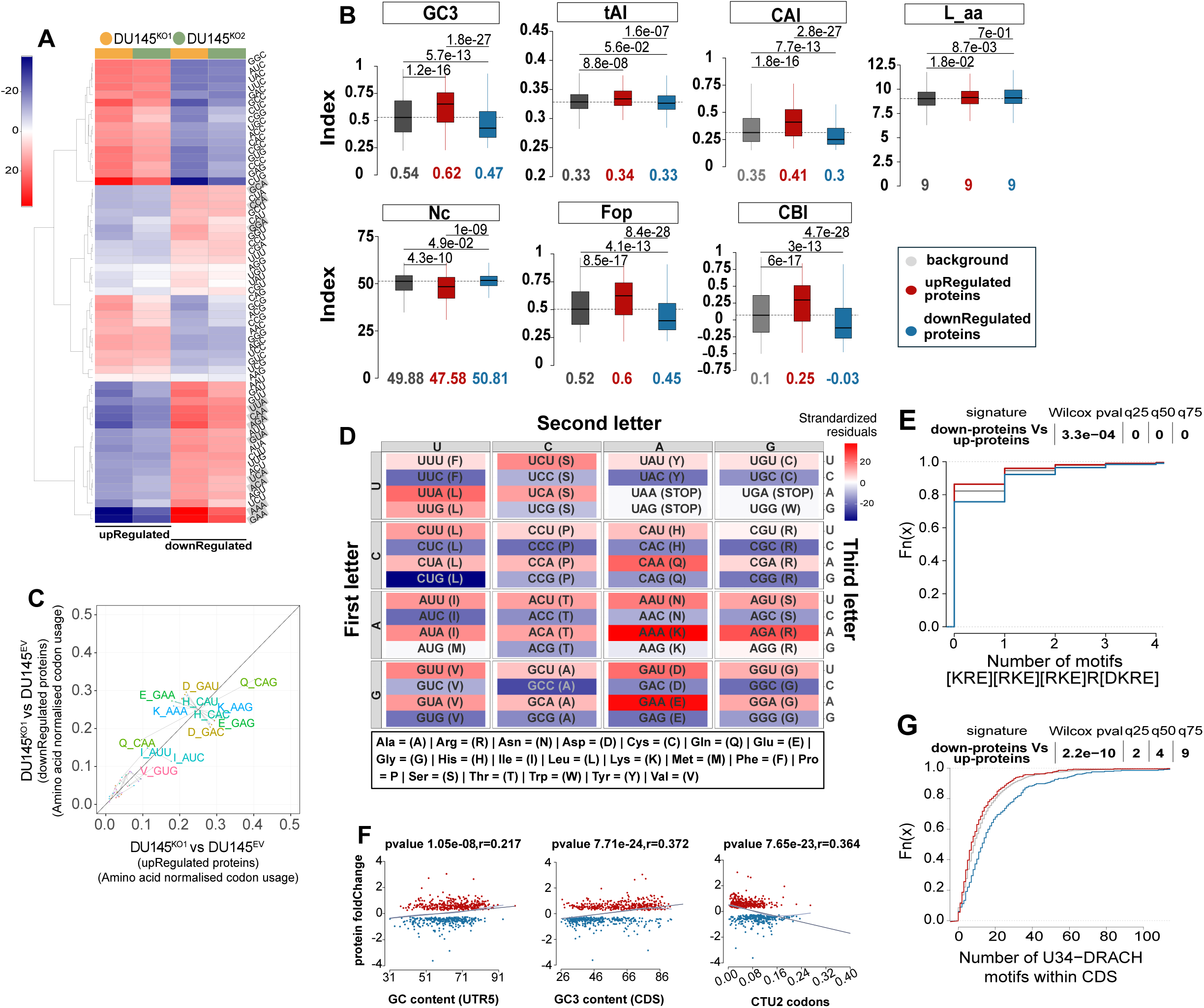
Coding sequence features associating with ELP3-sensitive protein expression. A) Heatmap displaying standardized residuals from chi-squared contingency tests. Red indicates codons overrepresented, and blue codons underrepresented, relative to the expected frequencies assuming independence between codon usage and protein regulation. U34-decoded codons are highlighted in gray, and codons are grouped by unsupervised hierarchical clustering. B) Boxplots comparing codon usage metrics between mRNAs encoding regulated proteins and the background set (gray). Seven measures were assessed: GC content at the third codon position (GC3s), tRNA Adaptation Index (tAI), Codon Adaptation Index (CAI), Effective number of codons (Nc), Frequency of optimal codons (Fop), Codon Bias Index (CBI), and number of amino acids (L_aa). Median values are indicated below each boxplot, and statistical significance was evaluated using a two-sided Wilcoxon rank-sum test. Median values are indicated below each boxplot. C) Comparison of codon usage, normalized to amino acid composition, between proteins downregulated and upregulated upon ELP3 loss (DU145^KO1^ Vs. DU145^EV^). Codons for the same amino acid are grouped by gray lines and depicted in matching colors. D) Codon table view of standardized residuals from chi-squared contingency tests for down-regulated proteins. Rows and columns correspond to the first, second, and third nucleotide positions, respectively. Cell colors reflect residuals, with positive values indicating enrichment and negative values indicating depletion relative to the null expectation. Amino acid one-letter abbreviations are explained in the reference box below the codon table. E) Empirical cumulative distribution functions (ECDF) of pentamer hydrophilic motif enrichment. Distributions are shown for background (gray), down-regulated proteins (blue), and up-regulated proteins (red). Statistical significance of the difference between up- and down-regulated groups was assessed by a two-sided Wilcoxon rank-sum test. P-values and the magnitude of distribution shifts at the 25th, 50th, and 75th percentiles are reported. F) Scatterplots showing selected features from Model 1 showed in Fig. 3B-C (left to right: 5′UTR GC%, CDS GC3%, and CTU2-dependent codon frequency) plotted against log₂ fold changes of significantly regulated proteins upon ELP3 suppression. Pearson correlation coefficients and corresponding p-values are indicated. G) Same analysis as in (E), but applied to DRACH motifs containing U34-tRNA decoded codons.

**extendedFigure3_02:**
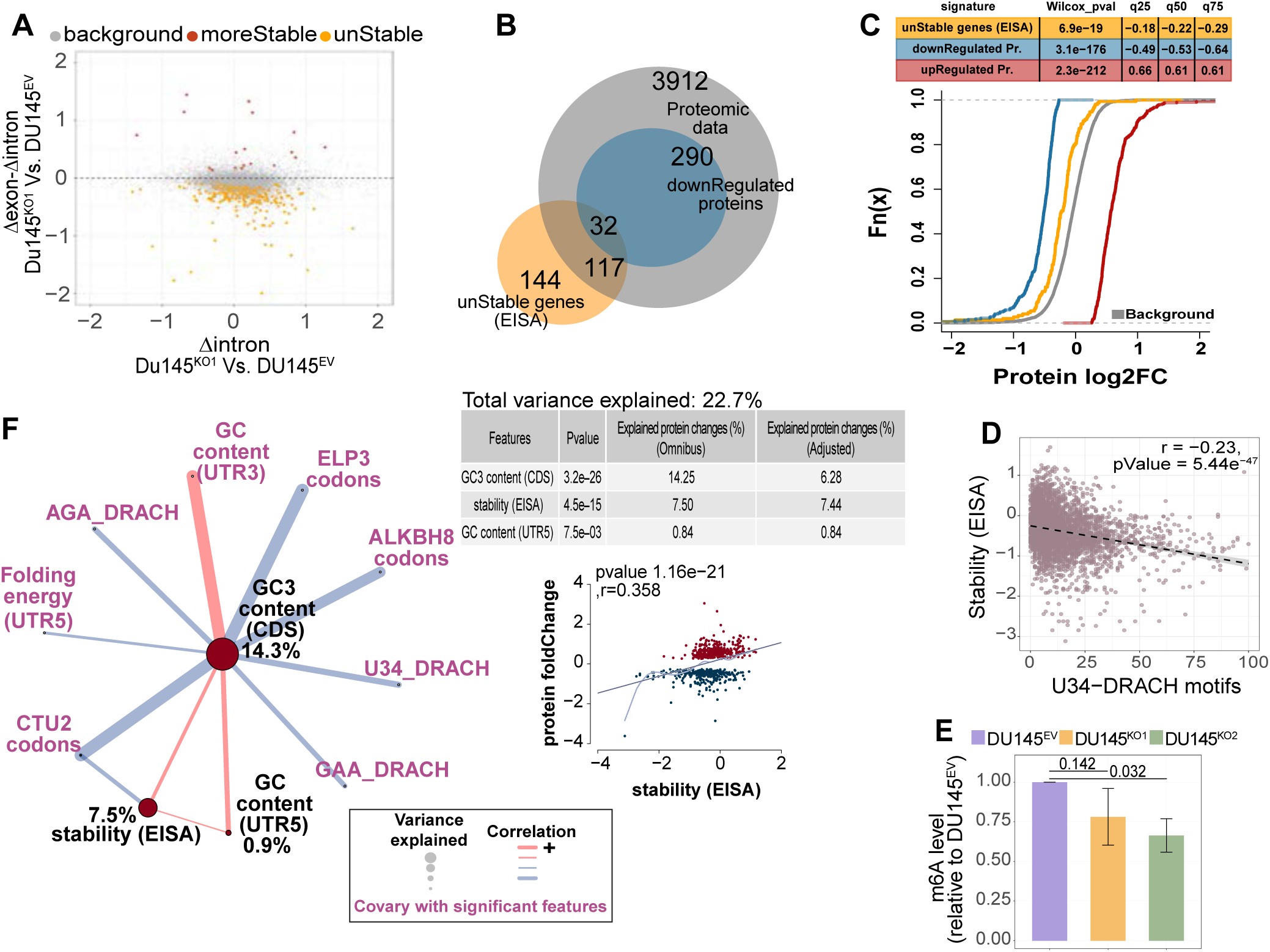
ELP3-sensitive RNA stability associates with frequency of U34-DRACH motifs. A) Exon–intron split analysis (EISA) using RNAseq data from DU145^KO1^ and DU145^EV^ cells. The x-axis shows intronic log2 fold-changes (Δintron, reflecting transcriptional input), while the y-axis shows exon–intron log2 differences (Δexon – Δintron, reflecting post-transcriptional regulation, i.e., RNA-stability). Each point represents one transcript. Transcripts classified as more stable are highlighted in red, unstable transcripts in orange, and background in gray (FDR < 0.05, quasi-likelihood framework (QLF)). B) Venn-diagram illustrating the overlap between unstable transcripts identified by EISA and downregulated proteins detected by proteomics. (gray circle = all quantified proteins) C) Empirical cumulative distribution functions (ECDFs) of protein log₂ fold changes between DU145^KO1^ and DU145^EV^ cells. Unstable transcripts identified by EISA are shown in orange, alongside down- and upregulated protein sets, compared to the background (gray). Significant shifts relative to the background were assessed using a two-sided Wilcoxon rank-sum test, reporting p-values and shifts at the 25th, 50th, and 75th percentiles. D) Scatterplot showing the relationship between U34-dependent DRACH motif counts and EISA stability scores (signed –log10 FDR values, reflecting both significance and direction of exon–intron changes) across all proteins detected by proteomics. E) m□A levels in poly(A)+ RNA isolated from DU145^EV^, DU145^KO1^, and DU145^KO2^ cells, shown relative to DU145^EV^. Bars represent mean ± SD from 3 biological replicates. Statistical significance was assessed by one-way ANOVA followed by Tukey’s HSD post-hoc test, with adjusted p-values shown. F) Network representation of Model 1 including EISA stability scores as an additional feature. Statistics and percentage of explained protein-level regulation is summarized for selected features (top right). The correlation between stability scores and proteomic fold changes is displayed as a scatterplot with Pearson correlation and p-value (bottom right).

**extendedFigure3_03:**
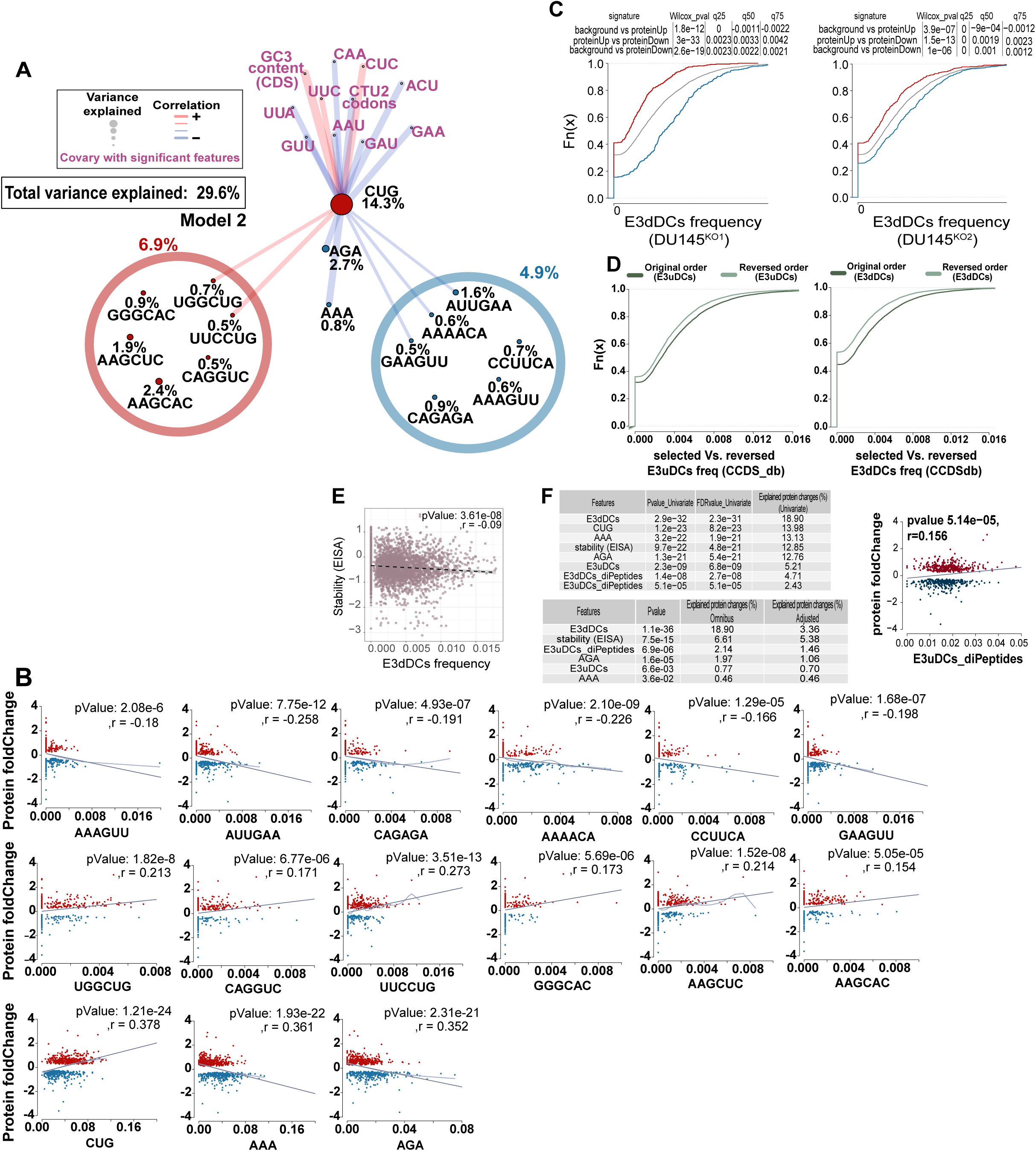
Di-codon context, rather than single-codon usage, accounts for proteomic changes upon ELP3 suppression. A) Network representation of Model 2 (Fig. 3B) result. Shown are the percentage of proteomic regulation explained by the model with additional breakdown by feature category. Features labeled in black collectively contribute to the explained variance in the model while purple features co-vary. Edge thickness represents the correlation strength between features, while node colors denote associations with protein down- or up-regulation upon ELP3 loss. B) Scatterplots of Model 2 selected features versus log₂ fold changes of significantly regulated proteins following ELP3 loss. Pearson correlations and associated p-values are shown. C) ECDFs of E3dDC frequencies in transcripts corresponding to upregulated (red), downregulated (blue), and background (gray) proteins. Results are shown separately for DU145^KO1^ (left) and DU145^KO2^ (right). Wilcoxon p-values and distribution shifts at the 25th, 50th, and 75th percentiles are also indicated. D) Empirical cumulative distribution functions (ECDFs) of di-codon frequencies in the CCDS database. Left: E3uDCs (di-codons linked to protein up-regulation) in their original versus reversed order. Right: E3dDCs (di-codons linked to protein down-regulation) in their original versus reversed order. E) Scatterplot depicting the correlation between E3dDC frequency and EISA stability scores (signed –log10 FDR values, reflecting significance and direction of exon–intron changes) across all proteins detected by proteomics. F) (Left) Selected features from the extended model (Fig. 3F) are summarized in two tables. The top table reports p-values, adjusted p-values, and percentage of regulation explained in univariate models, whereas the bottom table shows omnibus p-values, variance explained, and adjusted variance after accounting for all other selected features in the multivariate model. (Right) Scatterplot depicting the correlation between E3uDCs-dipeptide frequency and proteomic fold changes, with Pearson correlation statistics indicated.

**extendedFigure4_01:**
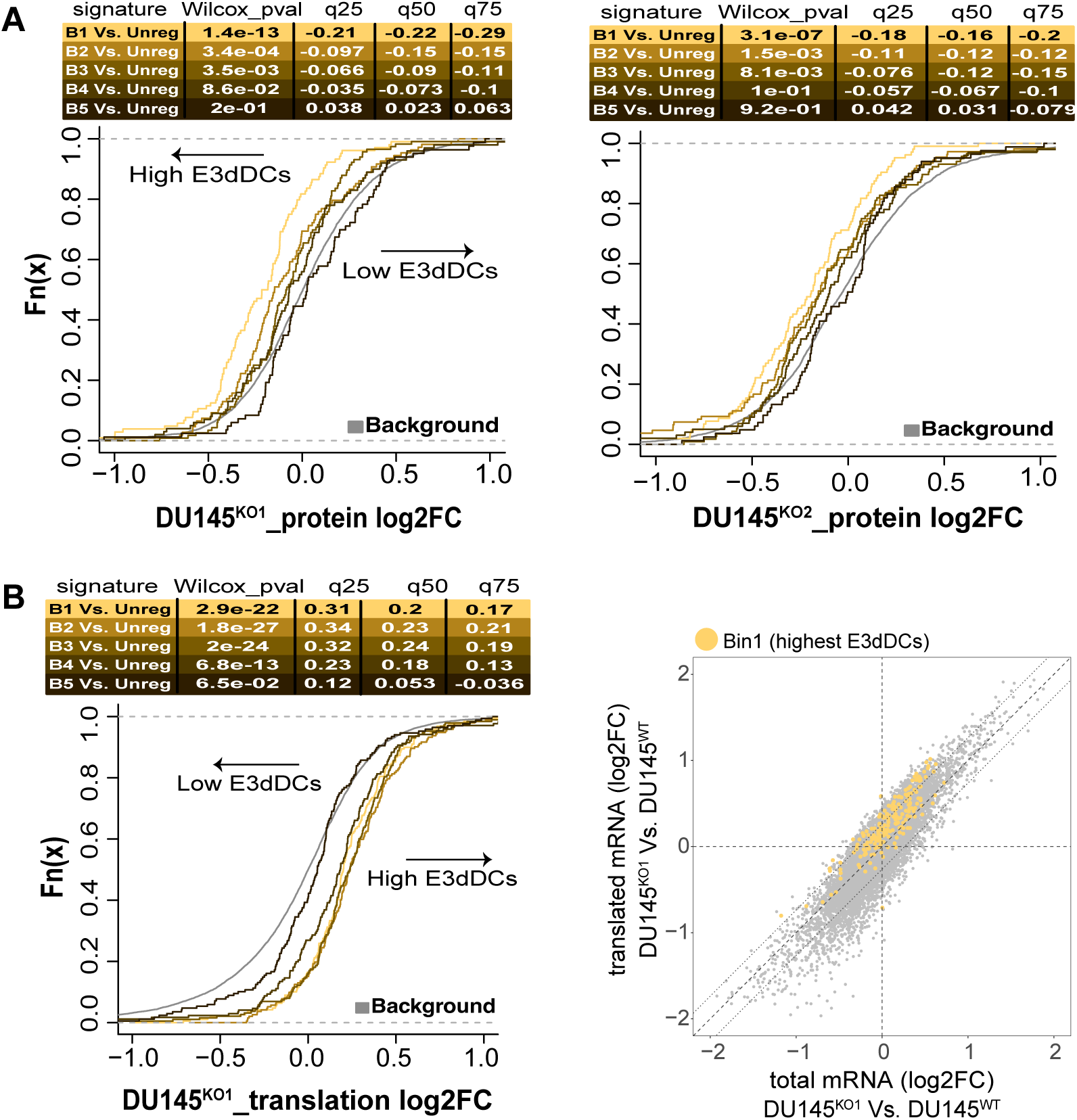
E3dDC-associated ISR failure in triggering selective mRNA translation upon ELP3 suppression. A) Empirical cumulative distribution functions (ECDFs) of protein abundance changes for ISR signature transcripts^30^, stratified into bins according to E3dDC frequency (bin 1 = highest, bin 5 = lowest). Non-regulated proteins are considered background (gray). Significant shifts relative to background were assessed using Wilcoxon rank-sum tests, with p-values and quartile (25–75%) shifts indicated. Left-shifted curves correspond to down-regulation upon ELP3 suppression. Left: DU145^KO1^ proteome; Right: DU145^KO2^ proteome. B) (Left) Same analysis as in (A), but showing ECDFs of log₂ fold-changes in translation (i.e. modulation of polysome-associated mRNA adjusted for total mRNA alterations) from anota2seq analysis of DU145^KO1^ Vs. DU145^EV^. Right-shifted curves correspond to increased polysome association upon ELP3 suppression. (Right) Scatterplot from anota2seq translatome analysis (DU145^KO1^ Vs. DU145^EV^), highlighting mRNAs from the ISR signature subset most enriched in E3dDCs.

**extendedFigure5_01:**
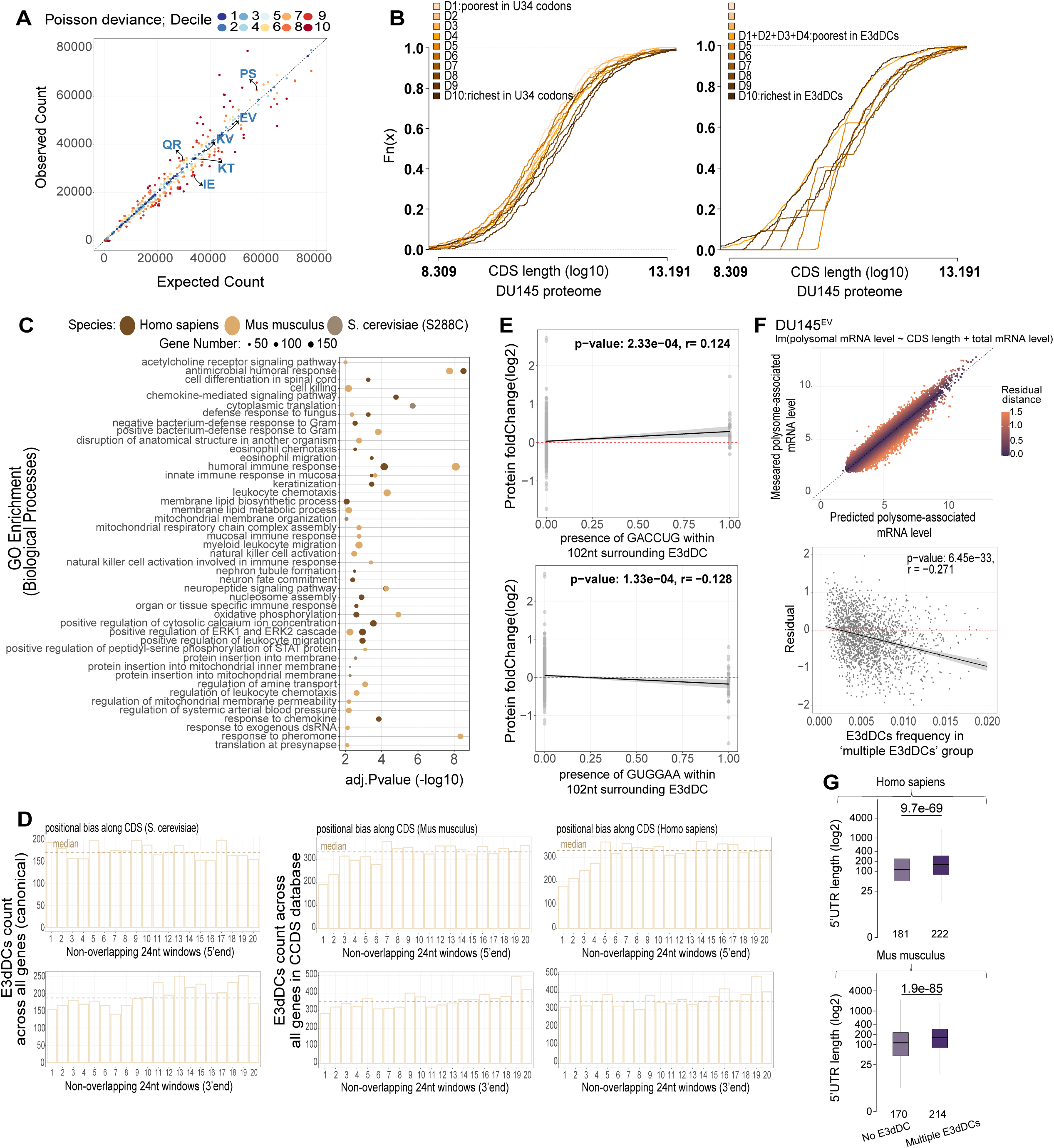
Cumulative presence and spatial distribution of E3dDC shape mRNA sensitivity to ELP3-dependent translational defects. A) Scatterplot comparing observed and expected di-peptide counts from the longest CDS isoforms in the CCDS database. Expected values were derived from a zero-order Markov model, which predicts di-peptide frequencies from the product of independent single–amino acid frequencies, assuming no positional dependence. Each point corresponds to a di-peptide, colored by Poisson deviance decile (dark blue = lowest, dark red = highest). The diagonal denotes equality between observed and expected counts. Di-peptides corresponding to E3dDCs are indicated. B) Empirical cumulative distribution functions (ECDFs) of CDS length (log₁₀) across the DU145 proteome. (Left) Proteins were stratified into deciles based on U34-codon count, from D1 (lowest) to D10 (highest). (Right) Proteins were stratified into deciles according to E3dDC content, with D1–D4 representing no E3dDCs while D10 the bin with the highest E3dDC count. C) Cross-species GO enrichment analysis of transcripts lacking E3dDCs in their CDS. Displayed are biological process (BP ontology) terms that were considered significant (FDR<0.01) in at least one of the three species (human, mouse, yeast). Dot size reflects the number of genes per species, and color indicates species origin. The x-axis shows enrichment significance. D) Positional distribution of E3dDCs across all transcripts in yeast (*S. cerevisiae*), mouse, and human (left to right). Coding sequences were partitioned into consecutive, non-overlapping 24-nt windows from the 5′ (top) and 3′ (bottom) ends. Bars indicate the total number of E3dDCs across transcripts within each window, with the line showing the median count across windows. E) Local sequence features surrounding E3dDC influence protein abundance changes in proteins with a single E3dDC. Scatterplots show protein log₂ fold-changes as a function of the presence of a specific di-codon within 102 nt around the E3dDC. Presence of GACCUG (top) is associated with increased protein abundance, while GUGGAA (bottom) is linked to decreased abundance (Pearson’s r and p-values shown; model FDR < 0.05). F) Linear regression model of polysome-associated mRNA levels in DU145^EV^. (Top) Measured versus predicted values from a model incorporating CDS length and total mRNA abundance as predictors. The color scale indicates residual distance from the regression fit. (Bottom) Residuals are plotted against the frequency of E3dDCs in the multiple di-codon protein group, with Pearson’s r and p-value indicated. G) Boxplots comparing 5′UTR length between mRNAs with multiple E3dDCs in their coding region and those lacking E3dDCs. Analyses were performed separately for human (top) and mouse (bottom). Mean lengths (unlogged) are shown below each boxplot, and p-values from Wilcoxon rank-sum tests are indicated.

**extendedFigure5_02:**
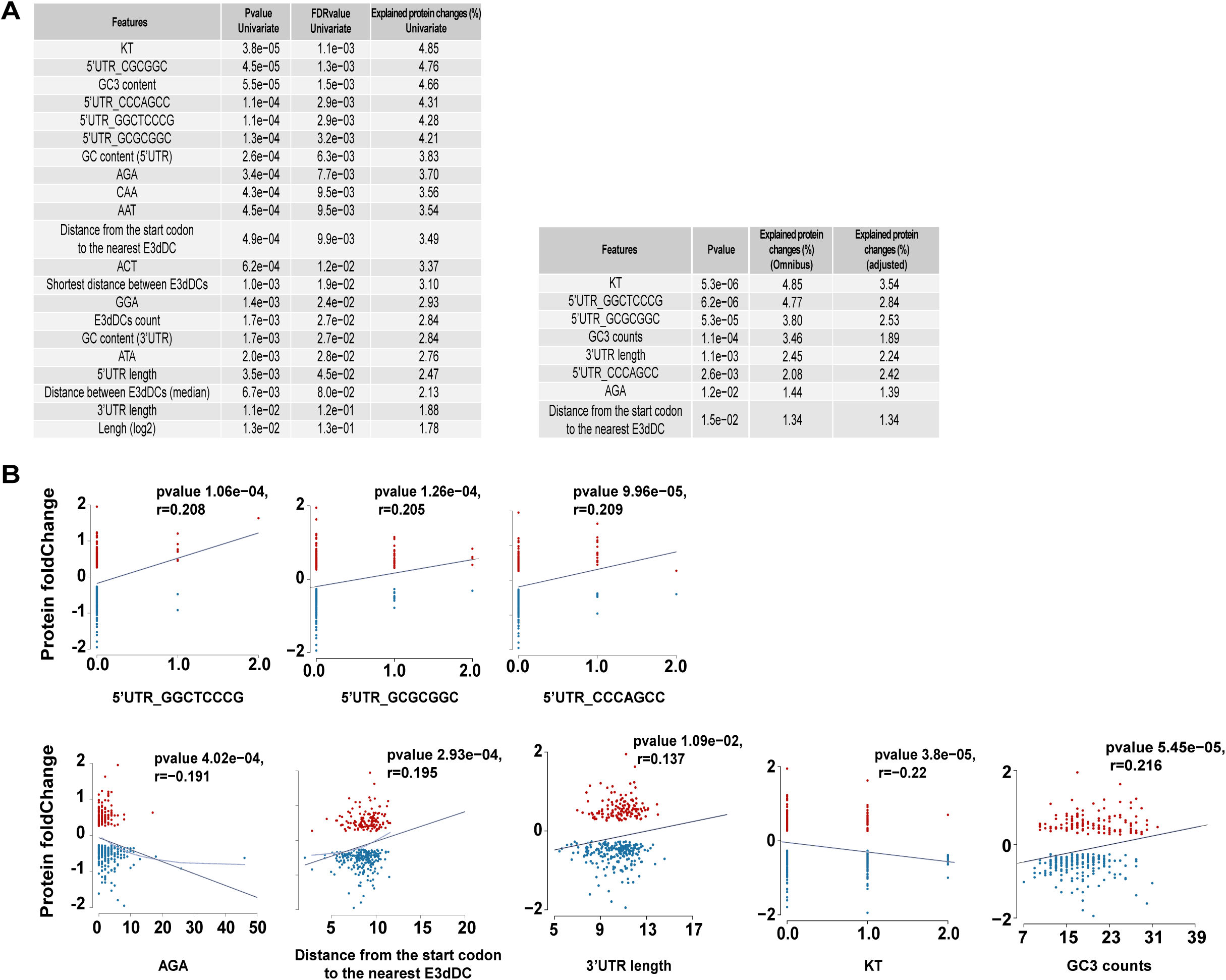
E3dDC context associated with ELP3-dependent translational defects. A) Features selected during postNet analysis within the subgroup with multiple E3dDCs are summarized in two tables: the left displays p-values, adjusted p-values, and variance explained from univariate models, while the right presents omnibus p-values, variance explained, and adjusted variance after adjusting for all other selected features in the multivariate model. B) Scatterplots of postNet-selected local features, including number of RBP motifs, AGA codon count, distance from the start codon to the nearest E3dDC, length of 3’UTR, KT di-peptide counts and GC3 content, plotted relative to log₂ fold-changes of significantly regulated proteins with multiple E3dDCs following ELP3 suppression. Pearson’s r and corresponding p-values are indicated.

**extendedFigure6_01:**
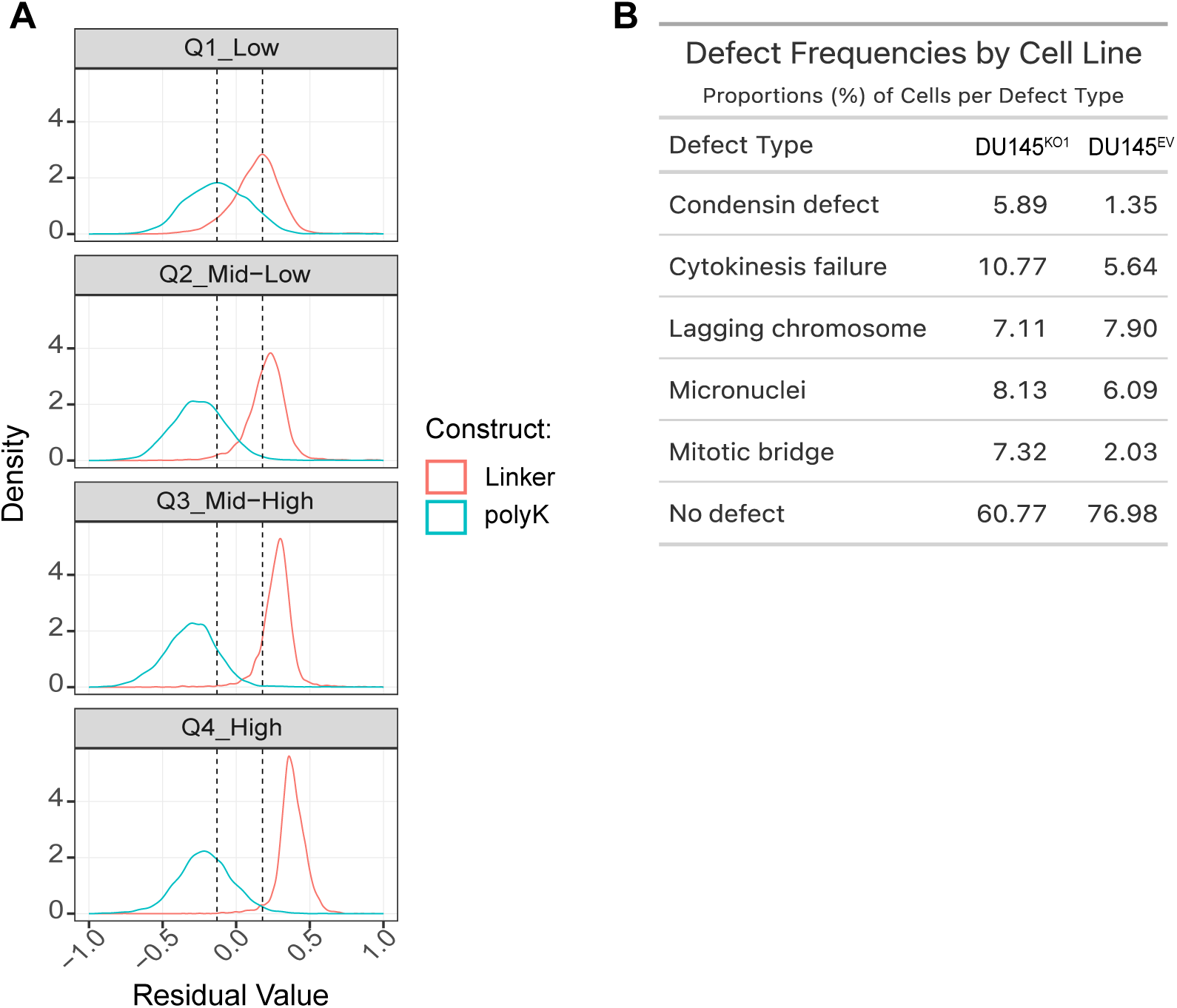
Translation reporter-assay quality-control and quantification of mitotic defects. A) Densities of regression residuals (mCherry ∼ GFP + replicate + cellType) from cells expressing linker (red, efficient translation) or polyK (cyan, inefficient translation) constructs, stratified into GFP intensity quartiles (Q1–Q4). The separation between linker- and polyK-expressing cells increases progressively from low (Q1) to high (Q4) GFP intensities, with the strongest differences observed in the upper two quartiles. This is consistent with a wider dynamic range in cells expressing higher levels of the reporters. B) Quantification of mitotic defects in DU145 cells. Frequencies of individual defect types (condensin defects, cytokinesis failure, lagging chromosome, micronuclei, mitotic bridge) are shown as proportions of cells in DU145^KO1^ and DU145^EV^. The proportion of cells without detectable defects (“No-defect”) is also indicated.

